# Adaptation and conservation of turtle visual opsins over a deep phylogenetic divergence

**DOI:** 10.64898/2026.10.31.763363

**Authors:** Taegan JM Perez, Golnar Jalilvand, Wali Mir, Angela Adjei, Belinda SW Chang, Rayna C Bell, Ryan K Schott

**Affiliations:** Department of Biology & Centre for Vision Research, York University, Toronto, Canada; Herpetology Department, California Academy of Sciences, San Francisco, CA, United States of America; Departments of Ecology and Evolutionary Biology & Cell and Systems Biology, University of Toronto, Toronto, ON M5S 3G5, Canada; Centre for the Analysis of Genome Evolution and Function, University of Toronto, Toronto, ON M5S 3B2, Canada

**Keywords:** Retinal photoreceptor cells, visual pigments, visual opsin, phototransduction, ultraviolet sensitive

## Abstract

The ecological radiation of organisms in diverse light environments depends, in part, on adaptive alterations within visual systems, particularly retinal photoreceptors and their visual pigments. Evidence of visual adaptation occurs across vertebrate lineages but is notably lacking in turtles, despite their complex visual system with up to eight photoreceptor cell types and five visual pigments. This lack of knowledge is especially pronounced in side-neck turtles (Pleurodira), as nearly all information on turtle vision derives from hidden-neck turtles (Cryptodira). Here, we bridge this ∼200-million-year evolutionary gap by investigating visual opsin evolution using an eye transcriptome and whole-genome sequences from 59 turtle species spanning both suborders, as well as an additional 42 representative reptile species. We estimated selective pressures using codon-based evolutionary models, testing for positive selection and long-term shifts between turtles and other reptiles, between suborders, and among major habitat types. We also analysed spectral tuning sites to assess how spectral sensitivity diverged between and within suborders. Our analyses revealed remarkably conserved turtle visual opsin sequences, with phylogeny-wide positive selection detected only in *LWS*. Habitat explained more shifts in selective pressure than taxonomy, with long-term shifts detected across all five opsins. Spectral tuning sites predict conserved UV sensitivity of *SWS1* across cryptodirans, consistent with the few species previously characterised, whereas shared changes identified in pleurodirans predict violet sensitivity, revealing previously unrecognized divergence in short-wavelength vision between turtle suborders. This study advances our understanding of turtle visual system evolution and highlights the need for conservation strategies that account for their distinct visual capabilities.

## Introduction

Visual perception, essential for many animals, allows the interpretation of colour, shape, size, and movement of objects in the environment, influencing responses to changing conditions (Davies et al. 2012; Nilsson 2021). Sensitivity of the visual system varies greatly among animals, in part due to differences in visual opsins, the proteins found in retinal photoreceptor cells. These opsins combine with a light-sensitive chromophore derived from vitamin A to produce visual pigments that are most efficient at absorbing specific wavelengths of light and thus differ in their peak absorbance (λ_max_). In vertebrates, the rod opsin visual pigment, *RH1*, is typically expressed in rod photoreceptor cells that function in dim-light conditions and is most sensitive to middle wavelengths (“green”, λ_max_ 460–530 nm). Cone visual pigments are typically expressed in cone photoreceptors that mediate bright-light vision and comprise four major classes: *SWS1* (“ultraviolet”, λ_max_ ∼355–375 nm, or “violet”, ∼385–455 nm) and *SWS2* (“blue”, λ_max_ 400–470 nm), which are sensitive to short wavelengths; *RH2* (“green”, λ_max_ 470–530 nm), sensitive to middle wavelengths; and *LWS* (“red”, λ_max_ 500–620 nm), sensitive to middle-to-long wavelengths (Bowmaker 2008; Davies et al. 2012; Hauser et al. 2014; Yokoyama 2000; Corredor et al. 2022; Zhang et al. 2026). The vertebrate ancestor likely possessed one rod opsin (*RH1*) and four cone opsins (*SWS1*, *SWS2*, *RH2*, and *LWS*), but this ancestral repertoire has been extensively modified by gene loss and duplication across vertebrate evolution (Davies et al. 2012).

Variation in spectral sensitivity can arise from multiple molecular mechanisms, including differences in the type, number, and proportions of opsin genes expressed in different photoreceptors, specific amino acid changes in the opsin at “spectral tuning sites” that can shift λ_max_, and the presence of different types of chromophores (Parry and Bowmaker 2000). Vertebrates can use two types of chromophores: the more common is derived from vitamin A_1_ (retinal, A_1_), but some species can instead (or also) use a chromophore derived from vitamin A_2_ (11-*cis* 3,4-didehydroretinal, A_2_). The A_2_ chromophore is most often associated with the use of freshwater aquatic environments and produces a visual pigment that has a long-wavelength-shifted sensitivity, broader absorption spectra, reduced photosensitivity, and increased thermal noise relative to a visual pigment composed of the same opsin and the A_1_ chromophore (Donner 2020; Corbo 2021; Corredor et al. 2022). Additionally, in some vertebrates, oil droplets in cone inner segments can act as filters, narrowing the spectral range of light reaching the visual pigments, which can enhance colour discrimination and chromatic constancy (Liebman and Granda 1971; Vorobyev 2003; Corredor et al. 2022).

The substantial diversity in vertebrate visual sensitivity has evolved, at least in part, through adaptation to different light environments and visual demands (Davies et al. 2012; Hagen et al. 2023). Associations between ecological niche and visual pigment spectral tuning have been demonstrated across vertebrates (for reviews see Hunt et al. 2009; Davies et al. 2012; Simões and Gower 2017; Carleton et al. 2020; Donner and Yovanovich 2020). Functional adaptation can also involve non-spectral properties of visual pigments, including their activation and decay kinetics. For example, transitions from marine to freshwater habitats in teleost fishes involved substitutions in *RH1* that both red-shifted spectral sensitivity and accelerated pigment kinetics, which was hypothesized to enhance the speed of dim-light adaptation (Van Nynatten et al. 2021). Molecular evolutionary studies likewise reveal strong associations between habitat and behaviour and selective pressures on visual opsins across fishes, birds, mammals, amphibians, and non-avian reptiles (*e.g.*, Veilleux et al. 2013; Wu et al. 2016; Hauzman et al. 2017; Schott et al. 2019, 2024; Gutierrez et al. 2018; Gerwin et al. 2024). However, visual-system evolution may also retain a strong signature of shared evolutionary history. Across ray-finned fishes, accounting for phylogenetic relationships substantially reduced several apparent associations between ecology and spectral sensitivity, highlighting the importance of shared ancestry in shaping visual-system diversity (Schweikert et al., 2018). Similarly, opsin expression in blennies can show strong phylogenetic dependence despite transitions among aquatic, amphibious, and terrestrial habitats (Cortesi et al., 2026). Distinguishing adaptation to similar visual environments from the retention of ancestral visual-system characteristics therefore requires consideration of both ecology and evolutionary history. Turtles, however, remain conspicuously understudied in this broader comparative framework, despite early studies on their photoreceptor and retinal anatomy and function (e.g., Forbes et al. 1958; Northmore & Granda, 1991; Baylor and Hodgkin 1973; Kolb et al. 1987; Liebman and Granda 1971).

Turtles are comprised of two suborders, Pleurodira (side-necked turtles) and Cryptodira (hidden-necked turtles), which are most easily distinguished based on the way they retract their heads under the shell (Fig. 1; Thomson et al., 2021). Combined fossil and molecular evidence suggest that Pleurodira and Cryptodira diverged in the early Jurassic at approximately 199.5 million years ago (178.8–225.4 Ma; Shaffer 2009; Pereira et al. 2017). Only 93 of the 356 extant species of turtles are pleurodirans, and they are restricted to freshwater habitats, whereas some cryptodirans inhabit marine and fully terrestrial habitats (Gaffney et al. 2006; Lovich and Gibbons, 2021). Previous studies of turtle visual systems indicate that Cryptodira species in various aquatic habitats have different spectral sensitivities. Freshwater turtles, like the red-eared slider (*Trachemys* [*Pseudemys*] *scripta*), are most sensitive to longer wavelengths (orange, red) typical of the light environments in ponds, lakes, and rivers. By contrast, the strictly marine green turtle (*Chelonia mydas*), which inhabits clear, tropical waters, is broadly sensitive to blue, green, and yellow wavelengths (Granda and O’Shea, 1972). These species are also inferred to have different chromophores in their visual pigments (A_1_ in the green sea turtle and A_2_ in the freshwater slider) and have different colours and abundances of their accessory (oil droplet) pigments. Both the photopigment absorption and oil droplet complement contribute to the unique spectral sensitivities of each species (Liebman and Granda, 1971; Horch et al. 2008). In addition, behavioural colour discrimination extending into the ultraviolet (UV) and the presence of up to five single-cone types suggest the potential for tetrachromatic colour vision in cryptodiran turtles (Arnold & Neumeyer, 1987; Kolb & Jones, 1987; Emerling 2017). Despite these studies, our understanding of turtle vision remains limited and is primarily based on hidden-necked turtles (Cryptodira), leaving a significant gap in our understanding of side-necked turtles (Pleurodira) and, more broadly, visual evolution across the turtle tree of life.

**Figure 1.**
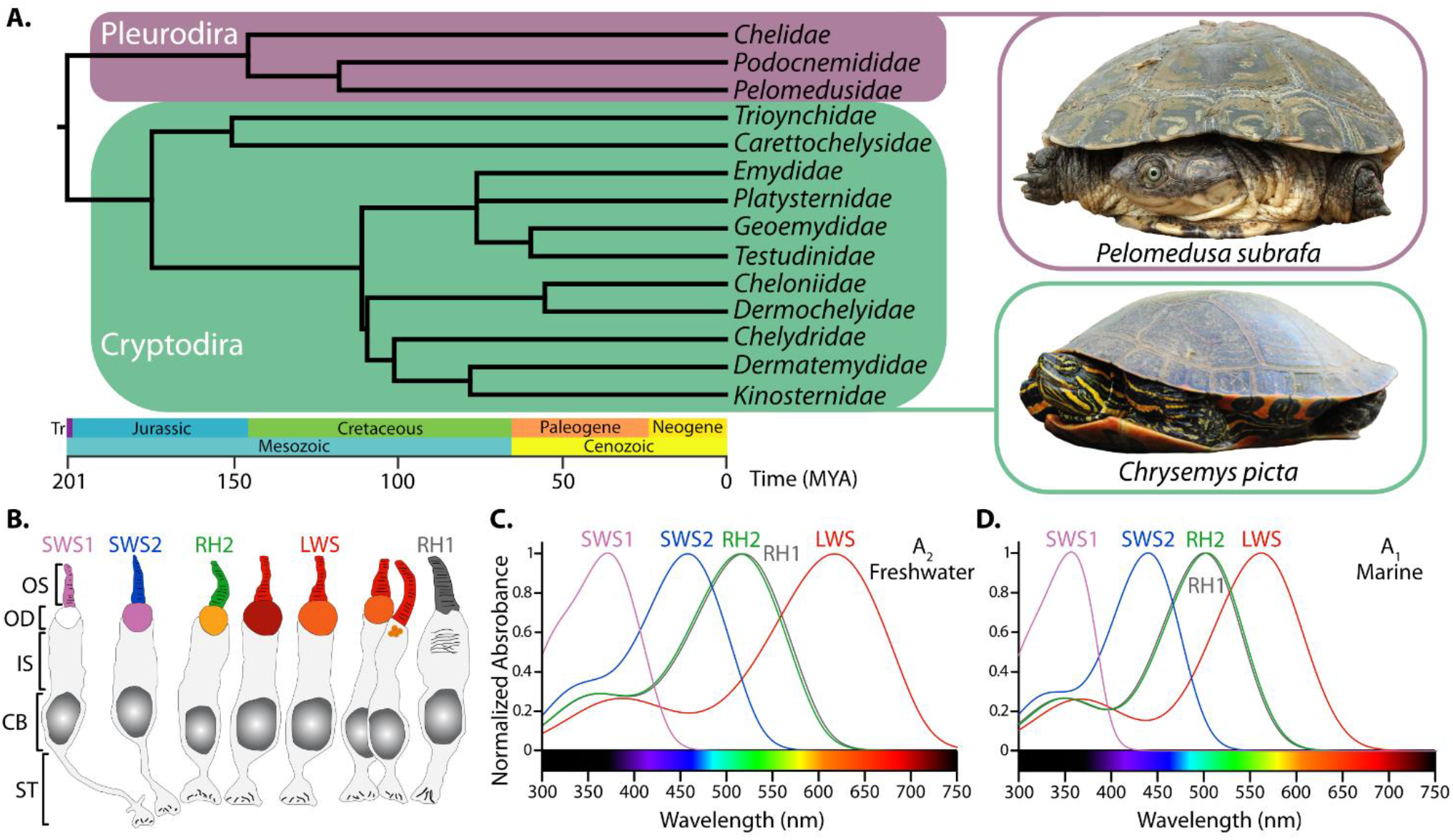
Overview of turtle phylogenetic relationships, divergence times, and photoreceptor systems. **A.**, Family-level phylogeny of turtles (Testudines) indicating the divergence time of the two suborders, Pleurodira (side-neck turtles) and Cryptodira (hidden-neck turtles) with examples of each taxon highlighting the difference in mode of neck retraction. **B.**, Illustration of turtle photoreceptors (*Trachemys scripta*), coloured based on the absorption spectra of the visual pigments in the outer segment and of the oil droplets. **C.** and **D.**, Normalized absorption spectra of a freshwater (*T. scripta*, **C.**) and marine (*Chelonia mydas*, **D.**) cryptodiran visual pigments. Freshwater cryptodirans tend to use the A_2_ chromophore resulting in broader, red-shifted spectra compared to the A_1_ chromophore used by marine turtles and most other marine and terrestrial vertebrates. Turtle phylogeny and divergence times based on Kumar et al. (2022). Photoreceptor illustrations based on Kolb (2013) and visual pigment and oil droplet absorbances based on Loew and Govardovskii (2001). Visual pigment absorption spectra are Govardovskii et al (2000) A_2_ and A_1_ template curves using maximal absorbances (λ_max_) estimated by MSP as follows: *Trachemys scripta* (Loew and Govardovskii 2001), *RH1* = 518 nm, LWS = 617 nm, *RH2* = 515 nm, SWS2 = 458 nm, SWS1 = 327 nm; *Chelonia mydas* (Liebman and Granda 1971), *RH1* = 502 nm, LWS = 562 nm, *RH2* = 502 nm, SWS2 = 440 nm. The *C. mydas* SWS1 λ_max_ of 358 nm was estimated from the amino acid sequence by Corredor et al. (2022). *Pelomedusa subrufa* and *Chrysemys picta* photographs by Johannes van Rooyen and Courtney Celley/USFWS, respectively. Abbreviations: OS, outer segments; OD, oil droplet; IS, innersegment; CB, cell body; ST, synaptic terminal.

Detailed retinal studies of the cryptodirans *C. mydas*, *Mauremys* (*Geoclemys*) *reevesii*, and *T. scripta elegans* have identified rods and multiple spectrally distinct single and double cones, including UV-, short-, middle-, and long-wavelength-sensitive cone classes associated with distinct oil droplets (Liebman and Granda 1971; Baylor and Hodgkin 1973; Ohtsuka & Kawamata 1990; Loew and Govardovskii 2001; Levenson et al. 2004; Horch et al. 2008; Grötzner et al. 2020). Based on their spectral sensitivities and comparisons with other vertebrates, these photoreceptors are predicted to express the ancestral complement of *RH1*, *SWS1*, *SWS2*, *RH2*, and *LWS*. However, although *RH1* expression in rods has been supported immunohistochemically, the molecular identities of the cone pigments remain incompletely characterised. Behavioural experiments nevertheless support tetrachromatic colour discrimination in several cryptodiran turtles (Arnold & Neumeyer 1987; Witherington & Bjorndal 1991; Kawamura et al. 2009). Whether this complex visual system is conserved across Cryptodira, and particularly across the unstudied Pleurodira, remains unknown.

The last common ancestor of extant turtles has been reconstructed as nocturnal, despite the predominance of diurnality among living species (Anderson and Wiens, 2017; Smith & Iverson, 2004; Hjort Toms et al., 2022; Chessman, 2024). Ancestrally nocturnal activity patterns, including hypothesized nocturnal bottlenecks, have profoundly shaped visual opsin evolution across vertebrates, including snakes (Castoe et al., 2013; Schott et al., 2018), geckos (Schott et al., 2019a), crocodilians (Emerling, 2016), and mammals (Wu et al., 2017). This contrast between reconstructed ancestral nocturnality and the complex cone visual system retained by, at least, living cryptodiran turtles provides alternative predictions for visual opsin evolution. If turtles experienced a prolonged nocturnal bottleneck, we predict reduced selective constraint and potentially loss of cone opsins. Conversely, if the complex cone system of extant cryptodirans reflects long-term maintenance of photopic vision, we predict retention of the ancestral vertebrate opsin repertoire under strong selective constraint and correspondingly strong conservation of opsin sequences and spectral tuning.

The deep evolutionary and ecological history of turtles provides an additional axis along which visual systems may have diversified. Turtles underwent early geographic and ecological expansion, including repeated transitions among terrestrial, freshwater, and marine habitats (Jaffe et al., 2011; Ferreira et al., 2015; Evers & Benson, 2019; Thomson et al., 2021). Although extant turtle diversity is low relative to many other reptiles, much of this ecological breadth persists among living lineages. The two turtle suborders nevertheless have markedly different evolutionary and ecological histories. Extant pleurodirans are less species-rich and ecologically diverse than cryptodirans, but their fossil record reveals substantial historical diversity, including extinct marine lineages (Gaffney et al., 2006; Ferreira et al., 2015; Evers & Benson, 2019). Their ∼200 My of independent evolution, coupled with ecological overlap among living species, therefore provides an opportunity to distinguish the relative influences of evolutionary history and ecology on visual opsin evolution. The deep divergence of Pleurodira and Cryptodira, together with ecological overlap among extant species provides an opportunity to assess whether visual opsin evolution more strongly reflects phylogenetic history or adaptation to visual environments. Greater differentiation between suborders would support a persistent influence of their independent evolutionary histories, whereas similarities among species occupying comparable habitats would point to ecological convergence. Strong conservation across turtle diversity would instead suggest substantial retention of ancestral visual opsin function.

Beyond phylogenetic differences, the ecological diversity of turtles provides several specific predictions for visual opsin evolution. Aquatic light environments vary strongly in spectral composition: marine waters are generally enriched in shorter wavelengths, whereas freshwater environments can be more spectrally variable and are often shifted toward longer wavelengths due to dissolved and suspended material (Costa et al., 2013; Hauser & Chang, 2017; Baden, 2024). Correspondingly, visual pigments of aquatic vertebrates often show habitat-associated spectral tuning, including shifts towards shorter wavelengths in marine environments and longer wavelengths in freshwater environments (Musilova et al., 2019; Van Nynatten et al., 2015, 2021). Although terrestrial habitats lack the strong spectral filtering of aquatic environments, terrestrial vertebrates also show visual-pigment adaptation to their photic environments (e.g., frogs (Schott et al., 2022, 2024) and mammals (Hunt et al., 2009; Gutierrez et al., 2018)). Given these patterns of visual opsin evolution across vertebrates, we predict habitat-associated differences in selective pressures and spectral tuning among marine, freshwater, and terrestrial turtles.

Marine turtles provide an additional opportunity to test for adaptation to rapid changes in light intensity. Some species routinely dive to depths where light becomes progressively dimmer and spectrally restricted, while morphological adaptations to marine locomotion permit more rapid movement through the water column than in terrestrial or freshwater turtles (Marvin & Lutterschmidt, 1997; Eckert, 2002; Blob et al., 2016; Hochscheid, 2014; Fossette et al., 2010; Van Nynatten et al., 2024; Jerlov, 1976). In some deep-diving vertebrates, including leatherback sea turtles, substitutions in *RH1* alter pigment kinetics, including accelerated retinal release rates that are hypothesized to facilitate rapid visual adaptation as animals move between bright surface waters and dim-light depths (Morrow and Chang, 2015; Van Nynatten et al., 2021; Xia et al. 2021; Dungan & Chang 2022). Whether other deep-diving sea turtles, or other visual opsins, show similar kinetic shifts has yet to be investigated. If the changes observed in leatherback *RH1* reflect broader evolutionary patterns in marine turtle vision, we predict distinct selective pressures on visual opsins, particularly at residues known to influence spectral tuning or pigment kinetics.

Marine turtles also encounter distinctive visual demands at night during reproduction. Females of many species emerge onto land to nest at night, while hatchlings typically emerge under low-light conditions and use UV visual cues for orientation towards the sea (Salmon et al. 1995; Witherington & Bjorndal 1991; Horch et al. 2008; Kawamura et al. 2009; Price et al. 2018; Vandersteen et al. 2020; Simantiris et al. 2025). The importance of short-wavelength and UV sensitivity during these critical life-history stages provides an additional basis for predicting distinct selective pressures on marine-turtle *SWS1*.

Here we produced the first eye transcriptome of a pleurodiran turtle and assembled visual opsin gene datasets using available genomes from 59 turtle species and an additional 42 representative reptile species. The genomic data allow us to investigate gene duplication and loss of the ancestral visual opsin repertoire across turtle diversity, while the eye transcriptome provides the first direct test of whether these opsins are expressed in the pleurodiran eye. We use codon-based models of molecular evolution to test for evidence of positive selection and long-term shifts in selective pressures in visual opsin genes between turtles and other reptiles, and among turtle species associated with phylogenetic and ecological transitions. We also investigate potential spectral tuning and changes at sites associated with visual pigment kinetics across these transitions. Finally, we test whether patterns of molecular evolution in turtle visual opsins are primarily explained by the ∼200-My divergence between Cryptodira and Pleurodira or by ecological transitions among freshwater, marine, and terrestrial habitats.

## Materials and Methods

### *Pelusios castaneus* Eye Transcriptome Sequencing and Assembly

A single eye was obtained from a *Pelusios castaneus* specimen accessioned at the University of Texas at Arlington Amphibian and Reptile Diversity Research Center. This specimen was initially identified as *Pelumedusa subrufa* but was later confirmed to be *Pelusios castaneus* through phylogenetic analysis. The eye was homogenized in Trizol (Invitrogen) using a BeadBug (Benchmark Scientific). Total RNA was extracted following a combined Trizol/RNeasy (Qiagen) protocol according to the manufacturer’s instructions. Library construction and sequencing on the Illumina HiSeq pipeline were performed according to standard protocols at The Centre for Applied Genomics, the Hospital for Sick Children (Toronto). Resulting 150-bp paired-end reads were trimmed using TrimGalore! (https://www.bioinformatics.babraham.ac.uk/projects/trim_galore/), which implements Cutadapt (Martin, 2011). Quality of processed reads was assessed with FastQC (http://www.bioinformatics.babraham.ac.uk/projects/fastqc/). Transcriptome assembly of each sample was performed *de novo* using Trinity (Grabherr et al. 2011) incorporating all paired reads following the standard protocol.

### Visual Opsin Gene Datasets

NCBI Genbank was searched using BLASTn for turtle visual opsin gene coding sequences (*RH1, RH2*, *LWS*, *SWS1*, *SWS2*) using queries obtained previously from other reptiles. Sequences from *Chrysemys picta* were used to assemble a reference set for subsequent searches of Genbank, the assembled *P. castaneus* eye transcriptome, and all available turtle genomes on the NCBI Genome database. BLASTn searches used the discontinuous megablast approach with an e-value cutoff of 1e-10. For the transcriptome searches, transcripts identified by BLASTn searches were imported into MEGA (Tamura *et al*., 2013), aligned with the reference, and manually trimmed to the coding sequence ensuring the longest open reading frame was recovered. For searches of genome assemblies, BLASTn results were parsed using a custom utility, and gene segments manually assembled, as needed, using MEGA and the corresponding reference sequence. An additional reference set of coding sequences was constructed from the *P. subrufa* data and used in subsequent rounds of BLAST searches. Visual opsin gene coding sequences for representatives of the other major reptilian lineages were obtained from NCBI BLASTn searches using queries from *Chrysemys picta* and *Sphenodon punctatus* following the method outlined above. A full list of species used for both analyses can be found in Supplementary File S1. We also searched the *P. castaneus* eye transcriptome for a complete coding sequence of the gene responsible for retinal chromophore switches *CYP27C1* using *Emydura macquarii* (XM_067570081) as a reference for BLAST searches as described above.

The recovered and assembled visual opsin gene coding sequences for each gene were first visually inspected in MEGA and aligned using MACSE (Ranwez et al., 2011) to produce in-frame codon alignments. To confirm accurate identification and alignment of all genes, Maximum Likelihood (ML) gene trees were inferred and inspected in MEGA using the GTR + G model before further analyses. Sequences with fewer than 50% of codons were excluded from molecular evolutionary analyses.

### Gene Synteny Analyses

In cases where a coding sequence could not be found, microsynteny analyses were conducted using the NCBI Comparative Genome Viewer (Rangwala et al., 2024). Orthologous genes flanking the target gene in an annotated genome from a closely related species were used to identify the corresponding genomic region in the target species. Regions extending 20 kb beyond the flanking genes were extracted from both species and aligned using minimap2 (-x asm10) and nucmer (--maxmatch; delta-filter -1). Coding sequences and individual exons were additionally searched against the corresponding genomic region and whole-genome assembly using BLASTn and tBLASTn, and candidate matches were evaluated to confirm opsin identity.

### Transcript Abundance Estimation

To qualitatively compare the relative abundance of visual opsins and other transcripts of interest, RNA-seq reads were quantified against a curated coding-sequence (CDS) reference rather than the *de novo* Trinity assembly, which contained fragmented and redundant contigs that could confound transcript-level abundance estimates. The reference included the five visual opsins, non-visual opsins, selected components of the rod and cone phototransduction cascades (*GRK1*, *GRK7*, *SAG*, *ARR3*, *GNAT1*, *GNAT2*, *PDE6B*, and *PDE6C*), retinal genes (*CRX*, *RLBP1*, and *RPE65*), genes associated with chromophore and carotenoid metabolism (*CYP27C1*, *CYP2J19*, *BDH1L*, and *TTC39B*) and broadly expressed reference genes (*RPL13A* and *GAPDH*).

*Pelusios castaneus* CDSs were extracted from the eye transcriptome using BLASTn searches, as described above, with sequences from the closely related turtle *Emydura macquarii* obtained from GenBank as queries. *Emydura macquarii* non-visual opsins were first identified by BLASTn searches of the NCBI core nucleotide database using frog non-visual opsins from Boyette et al. (2024) as queries. Similarly, *E. macquarii BDH1L* and *TTC39B* homologs were identified by BLASTn searches of the NCBI core nucleotide database using sequences from Toomey et al. (2022) as queries. For opsins not recovered from the *P. castaneus* eye transcriptome, the reference included *E. macquarii TMT3* and *Xenopus tropicalis* parietopsin and parapinopsin CDSs, the latter representing opsins not recovered in turtles. Inclusion of non-visual opsins provided alternative targets for reads derived from related opsin genes, reducing potential misassignment to the visual opsins. Similarly, to improve discrimination among closely related CYP paralogues, *CYP27A4* and *CYP2J2*-like CDSs were included as alternative mapping targets for *CYP27C1* and *CYP2J19*, respectively. These paralogues were identified by BLASTn as the sequences most similar to their respective focal genes and were included to reduce misassignment of reads from closely related CYP genes.

Transcript abundance was estimated using Salmon v2.8.0 (Patro et al. 2017) through the align_and_estimate_abundance.pl utility in Trinity v2.15.2, with GC-bias correction (--gcBias). The resulting abundance estimates were compiled into a transcripts-per-million (TPM) matrix using Trinity’s abundance_estimates_to_matrix.pl utility. Because only a single biological replicate was available, TPM values were used descriptively to compare relative transcript abundance, and no statistical comparisons of expression were performed.

### Molecular Evolutionary Analyses

Gene trees for each opsin (“gene tree”) were inferred by ML using PhyML 3 (Guindon et al. 2010) under the GTR+G+I model with a BioNJ starting tree, the best of NNI and SPR tree improvement, and aLRT SH-like branch support (Anisimova and Gascuel 2006). To produce “species tree” topologies for each gene, we generated a topology that matched expected species relationships in Reptilia modelled by TimeTree (Kumar et al., 2022) and trimmed the resulting phylogeny to match the taxon sampling of individual genes. For our analyses of turtles, we generated species topologies using a recent large-scale phylogeny (Thomson et al., 2021), again trimming these to match the sampling. The PAML 4 random sites models (M0, M1a, M2a and M8; (Yang 2007) using the BlastPhyMe software (Schott et al. 2019b) were employed to estimate the form and strength of selection acting on the visual opsin genes and to test for positive selection. As an additional test for episodic positive selection that accounts for potential synonymous rate variation the HYPHY model BUSTED (Murrell et al. 2015) was used as implemented on the Hyphlow platform and models were run in triplicate (Kwon et al., 2026). We also identified individual sites inferred to be under positive selection using the HYPHY model FUBAR (Murrell et al., 2013) on the same HYphlow platform. We used Clade model C (CmC) and Clade model D (CmD) to test for shifts in selection among the opsin genes in turtles compared to all other reptiles (Bielawski and Yang, 2004). We also performed a branch model test via PAML to test for differences in average selective constraint between turtles and other reptiles. We performed the same analyses using these models (except branch models) on the turtle only datasets. To test whether shifts in selection among opsin genes were primarily driven by phylogenetic divergence of Pleurodira and Cryptodira or shifts in habitat usage, we used CmC/CmD models. Specifically, we contrasted models that partitioned the two suborders with models that partition among the broad habitat categories of freshwater (which includes all Pleurodirans and a subset of Cryptodirans), marine, and terrestrial. We also used the HYPHY BUSTED model to test for episodic positive selection within these same partitions. Finally, we used the HYPHY model RELAX (Wertheim et al., 2015) on the HYphlow platform and ran in triplicate to test whether elevated ω values are likely to be the result of relaxed selective constraint (i.e., a lack of selection against change) or intensified selection (i.e., increased strength of negative and/or positive selection). A combination of likelihood ratio tests (LRTs) and Akaike Information Criterion (AIC) comparisons were used to determine the overall best-fitting model for each gene. Specifically, we determined the best-fitting model using the lowest AIC among CmC and CmD for both the gene and species tree topology analyses for each gene. We determined statistical support for the selected models using LRT against its corresponding null (M2a_rel for CmC; M3 for CmD).

We also identified variation at known spectral tuning sites and other functionally relevant sites between the two clades of turtles following (Schott et al. 2024). Bovine rhodopsin (*RH1*) (Palczewski et al. 2000) was used as a template. We compared these variant sites to those inferred to be under positive selection using the PAML and HYPHY models.

## Results

### All Five Visual Opsin Genes are Present in Both Side-neck and Hidden-neck Turtles

Coding sequences of the five visual opsin genes were extracted from NCBI GenBank, available genomes, and the *Pelusios castaneus* eye transcriptome, resulting in a dataset containing 59 turtle species. We found that all five genes are expressed in the *P. castaneus* eye transcriptome and present in the eight other pleurodiran genomes (nine including the *P. castaneus* genome; Fig. 2). Complete coding sequences of *RH1*, *RH2*, and *SWS1* were recovered from all species, whereas partial (>50% complete) or incomplete (<50%) sequences of *LWS* and *SWS2* were recovered from 13 species (partial: *LWS* = 8, *SWS2* = 7; incomplete: *LWS* = 3, *SWS2* = 1; Supplementary File S1). In all cases, recovered partial and incomplete sequences appeared otherwise conserved, with no evidence of sequence degradation or pseudogenization (*Zenodo dataset*).

**Figure 2.**
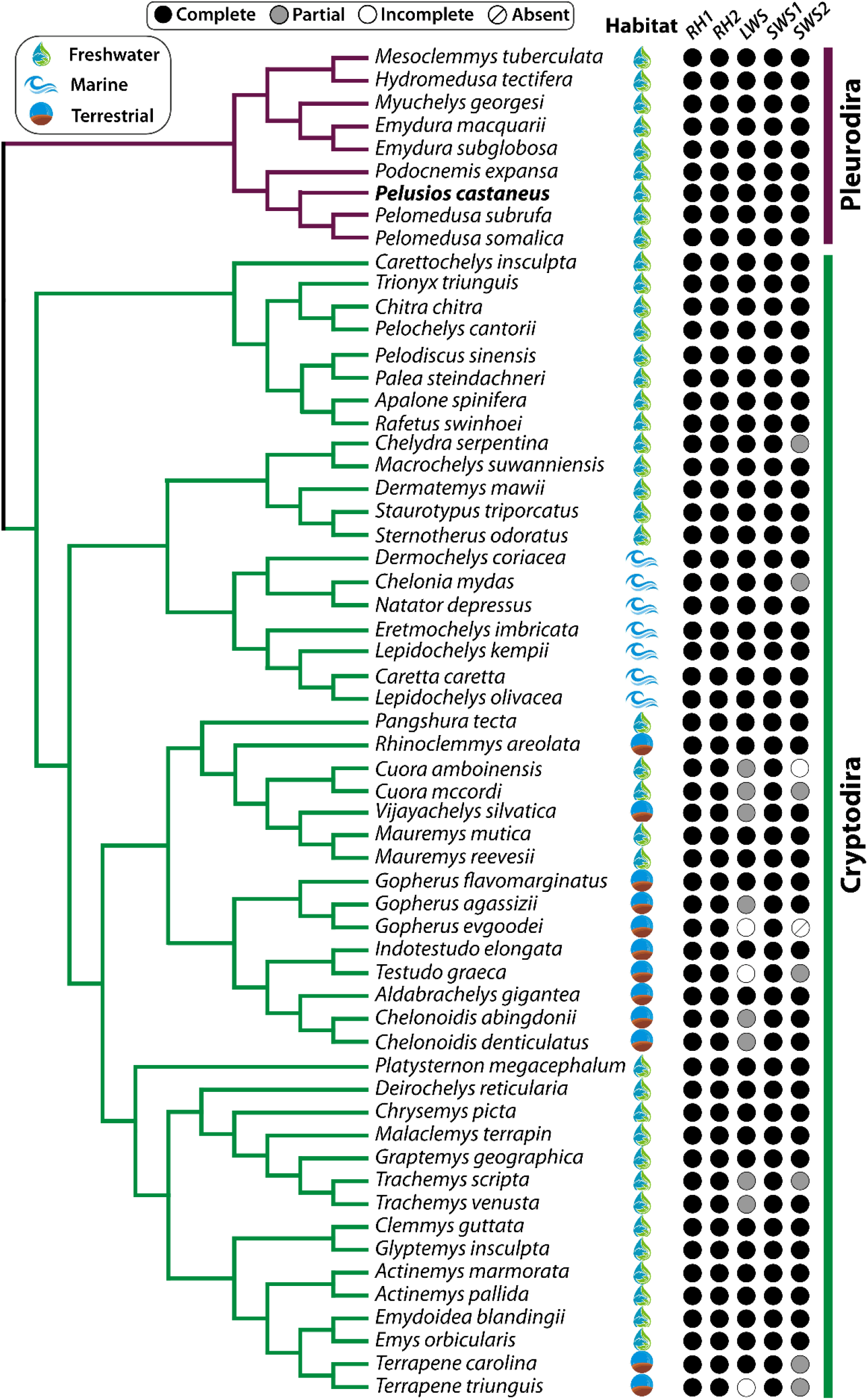
Species evolutionary relationships and recovery of turtle visual opsin gene sequences. Icons next to each species indicate their respective habitats: freshwater (droplet), marine (wave), terrestrial (circle). The recovery of each opsin coding sequence is also shown. Black circles indicate a complete sequence, grey circles a partial coding sequence (containing more than 50% of codons), and white (open) circles indicate an incomplete sequence (<50%), which was excluded from the molecular evolutionary analyses due to insufficient recovery. One sequence was completely absent indicated by the diagonal line through the open circle. The pleurodiran species *Pelusios castaneus* is represented by both genomic and transcriptomic data indicated by a bold species name. The topology is based on (Thomson et al. 2021).

Only a single species, *Gopherus evgoodei*, lacked a recoverable sequence entirely (*SWS2*), and this species also had an incomplete *LWS*. Because *LWS* and *SWS2* are adjacent in turtle genomes, we examined their expected genomic region in *G. evgoodei* using the congeneric *G. flavomarginatus* locus as a reference. The genes flanking the *LWS–SWS2* region were recovered in *G. evgoodei*, and whole-region alignments identified extensive homology between the two species across the broader locus (*Zenodo dataset*). However, the assembled *G. evgoodei* region was approximately 25 kb shorter than the homologous *G. flavomarginatus* region and contained a major alignment discontinuity within the opsin-containing interval. A 495-bp stretch of ambiguous nucleotides (Ns) occurred at the boundary of this discontinuity, immediately beyond the region containing recoverable *LWS* sequence. The annotation of this interval also differed between species: three long non-coding RNA loci occur between *LWS* and *SWS2* in *G. flavomarginatus*, whereas these loci were not annotated in *G. evgoodei*, which instead contained an intervening protein-coding locus, but one that lacked detectable homology to other annotated sequences across turtles. Targeted and genome-wide searches recovered sequences corresponding to the downstream portion of *LWS* but failed to identify *SWS2*-specific sequence. The status of *SWS2* in *G. evgoodei* remains unresolved, but the discontinuity and ambiguous sequence are consistent with the possibility of absence due to a sequencing or assembly artefact. More broadly, the restriction of partial, incomplete, and unrecovered sequences to the tandem *LWS* and *SWS2* loci, together with conservation of the recovered sequences and the absence of evidence for pseudogenization, is more consistent with difficulties in sequencing or assembling this genomic region than with repeated independent gene inactivation or loss across turtles.

### High Transcript Abundance of visual opsins and genes associated with chromophore and carotenoid metabolism

All five visual opsins were abundant in the eye transcriptome, with *LWS* (101,363 TPM) and *RH1* (88,994 TPM) showing the highest abundance, followed by *RH2* (31,437 TPM), *SWS2* (28,708 TPM), and *SWS1* (16,258 TPM; *Zenodo dataset*). Cone-associated phototransduction genes were also generally more abundant than their rod-associated counterparts: *GNAT2* (35,007 TPM) *vs GNAT1* (8,172 TPM), *PDE6C* (5,418 TPM) *vs PDE6B* (748 TPM), and *ARR3* (121,849 TPM) *vs SAG* (36,033 TPM), although *GRK7* (4,918 TPM) was slightly less abundant than *GRK1* (5,962 TPM). *CYP27C1* and *CYP2J19* were also abundant (7,367 and 5,203 TPM, respectively), at levels comparable to several phototransduction genes, while the additional carotenoid-associated genes *BDH1L* and *TTC39B* were expressed at 7,600 and 403 TPM, respectively. In contrast, the closely related alternative mapping targets *CYP27A4* and *CYP2J2*-like had lower abundances (828 and 585 TPM, respectively).

### Selective Constraint in Turtle Visual Opsins Compared to other Reptiles

Selection analyses were performed using “gene tree” (ML) and “species tree” topologies to test for robustness to topological differences. We used the PAML M0 model to determine overall selective constraint acting on visual opsins across reptiles, turtles, and non-turtle reptiles. Estimates of overall selective constraint were highly consistent between topologies, with M0 ω ranging from approximately 0.038 for *SWS1* to 0.103 for *RH1* across reptiles, and 0.046 to 0.143 in turtles (Fig. 3). Site-model comparisons provided no evidence of positive selection in any opsin under either topology in reptiles. However, with HYPHY BUSTED we found significant evidence for positive selection across reptiles in *LWS* (ω = 2.98 [1.09%], *P* = 0.01886) and *SWS1* (ω = 3.34 [0.42%], *P* = 0.0421; Supplementary File S2). Additionally, BUSTED analyses showed evidence of positive selection in *RH2* in non-turtle reptiles (ω_foreground_ = 83.131, ω_background_ = 0.744, *P* = 0.0413; Supplementary File S2). We used the PAML branch model to test for differences in average selective constraint between turtles and other reptiles and found evidence for significant and topology-robust differences for all the visual opsins, with greater selective constraint (lower ω) in turtle *RH2, LWS*, and *SWS2* (Fig. 3) and less constraint in *RH1* and *SWS1*. CmC likewise supported differences in selective pressure between turtles and other reptiles for all five opsins under both topologies, with the direction of the selection shifts matching those of the branch models (Fig. 3). Support for the more complex CmD was less consistent: *SWS1* strongly favoured CmD under both trees, whereas CmD support for *SWS2*, *LWS*, and *RH2* was sensitive to topology. *RH1* supported both clade models under both topologies, although their relative fit differed (Supplementary File S2). With RELAX we found evidence for a significant relaxation of selection in turtle *RH1* and *SWS1*, and intensification of selection in turtle *LWS* and *SWS2,* consistent with the patterns found using the branch and clade models with the exception of the lack of evidence in *RH2*.

**Figure 3.**
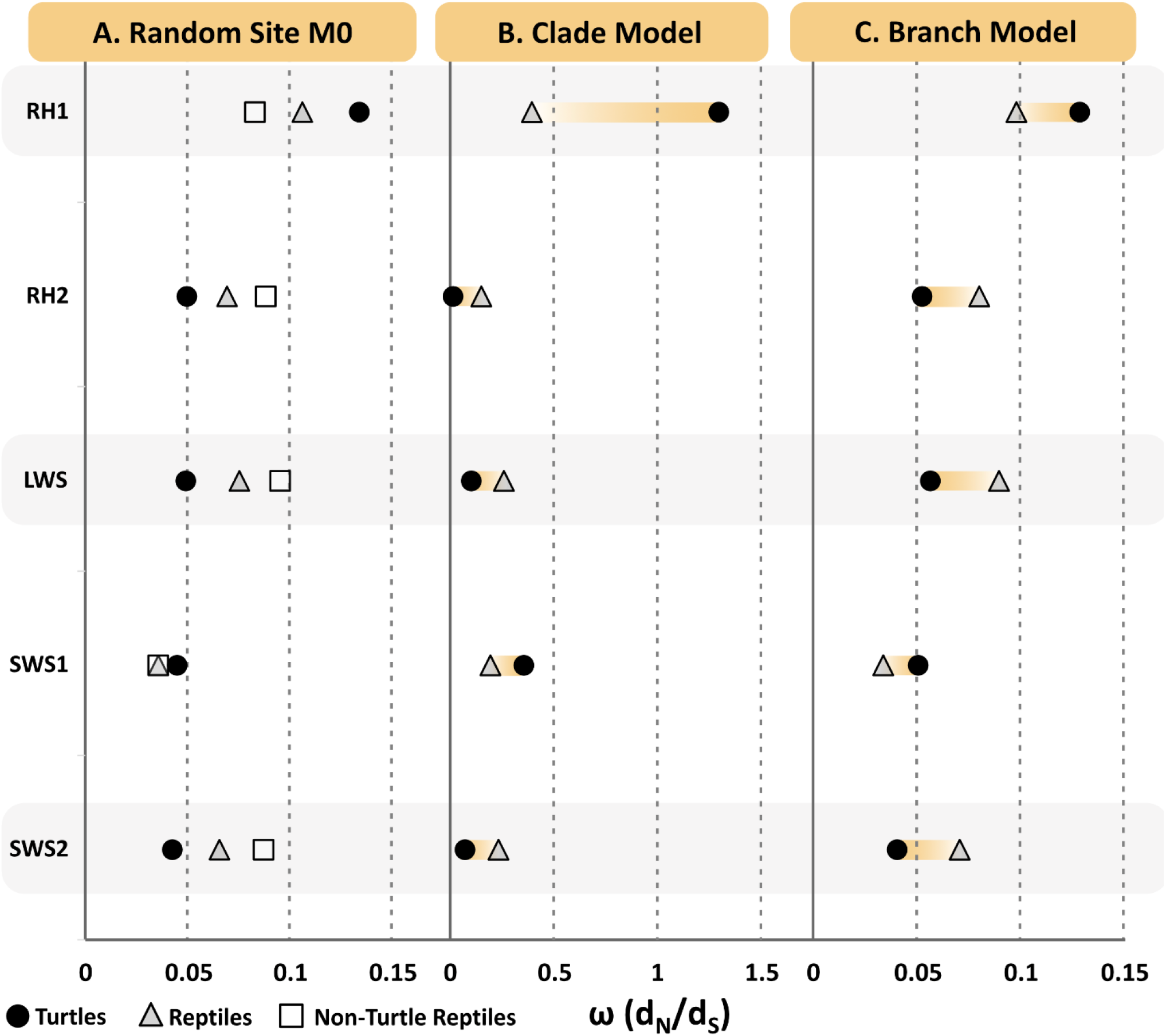
A. Selective pressures on reptile visual opsin genes estimated by PAML M0, Branch, and Clade models. **A.** PAML M0 results are shown for all reptile (triangles), non-turtle reptile (squares), and turtle-only (circles) datasets. The rod opsin, *RH1*, is under substantially less selective constraint compared to the four cone opsins genes (*RH2*, *LWS*, *SWS1*, *SWS2*) in both the all-reptile and turtle only tests. **B.** Results from best-fitting Clade model C or D showing evidence for differences in selective pressure between turtles and other reptiles. Values for turtles are shown with filled circles as the “foreground” while the remaining tested reptiles are shown with open circles as the “background”. **C.** Evidence for differences in selective constraint between turtles and other reptiles using PAML branch models.

### Shifts in Selective Pressures Among Turtle Opsin Genes

PAML clade model (CmC and CmD) analyses were used to compare selective pressures between Pleurodira and Cryptodira, as well as to test for evolutionary patterns in turtle opsins associated with phylogenetic history and habitat differences (specifically freshwater used by all pleurodirans and some cryptodirans vs marine and terrestrial habitats used by other cryptodirans). We found that CmC provided a better (or similar) fit for *RH1*, *RH2* and *LWS*, but that CmD fit better for *SWS1*, and *SWS2* (Supplementary File S2), presumably due to differences in the rate distribution profile and lack of support for a neutral site class (Van Nynatten et al., 2021). We report the better fitting of the two models below (see Supplementary File S2 for full results table). The marine *vs* non-marine partition was found to be the best-fitting partition for *RH1, RH2* and *LWS*, whereas the terrestrial *vs* aquatic partition was found to be the best fitting partition for *SWS1* and the freshwater partition for *SWS2*. For *RH1* and *LWS* the freshwater *vs* non-freshwater partition was also significant under CmC. For *LWS* the Pleurodira *vs* Cryptodira partition was also significant under CmD. For *SWS1* the freshwater *vs* non-freshwater partition was also significant under CmD. For *SWS2* the Pleurodira *vs* Cryptodira and Terrestrial partitions were also significant under CmD (Fig. 4).

**Figure 4.**
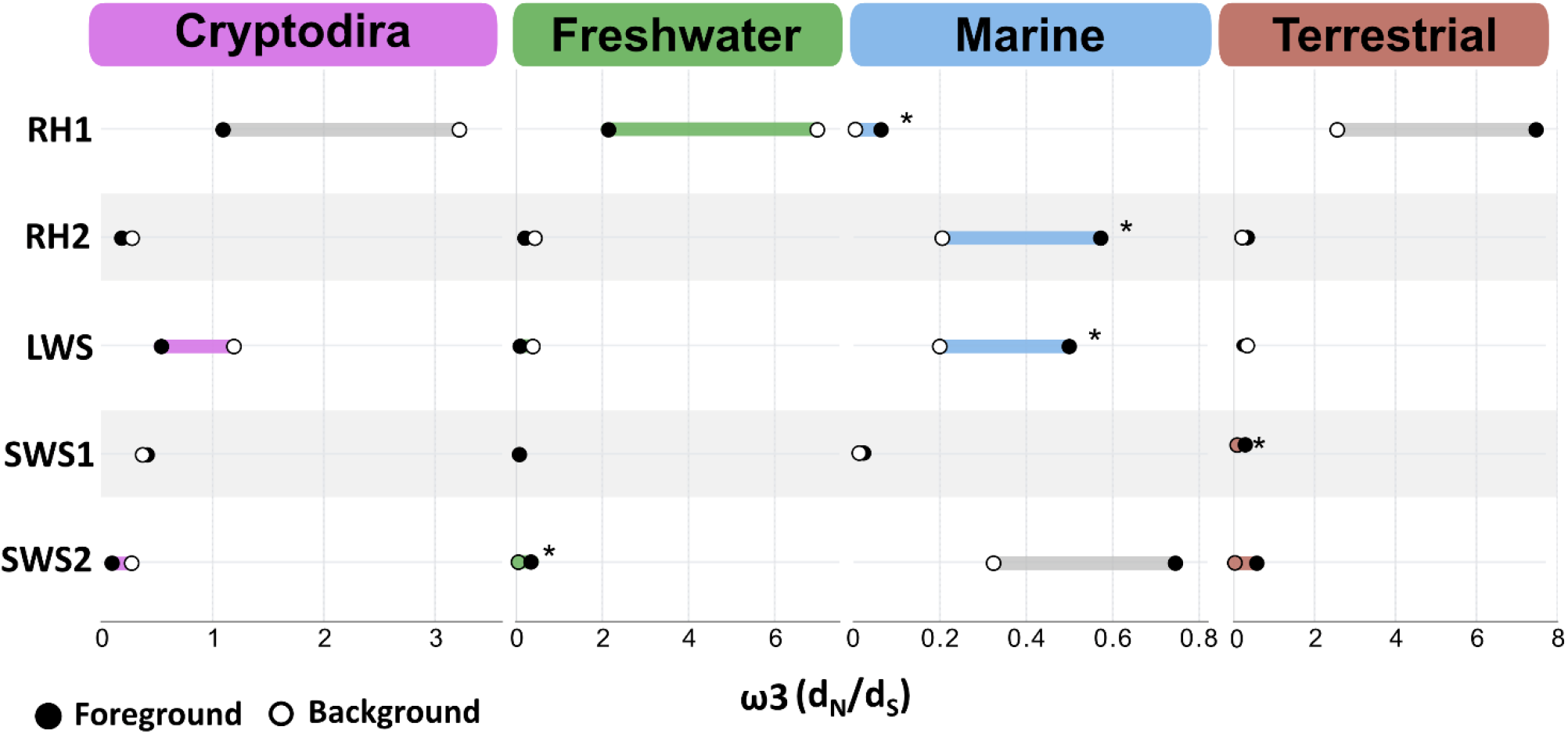
Selective pressures on turtle visual opsin genes estimated by PAML clade models (CmC and CmD). Estimated ω values for the divergent/third site class are shown for the foreground (closed circle) partition as named in each panel *vs* the background partition containing all other species (open circle). The better fitting of CmC and CmD and the gene (ML) and species tree topologies is shown for each partition (see Supplementary Table S2). Statistically significant results as determined by a Likelihood Ratio Test (LRT) to the appropriate null model (M2a_rel and M3, for CmC and CmD, respectively) are represented by coloured bars connecting the background and foreground ω values. The best-fitting partition for each gene is indicated with an asterisk. See Supplementary File S2 for full results.

**Figure 5.**
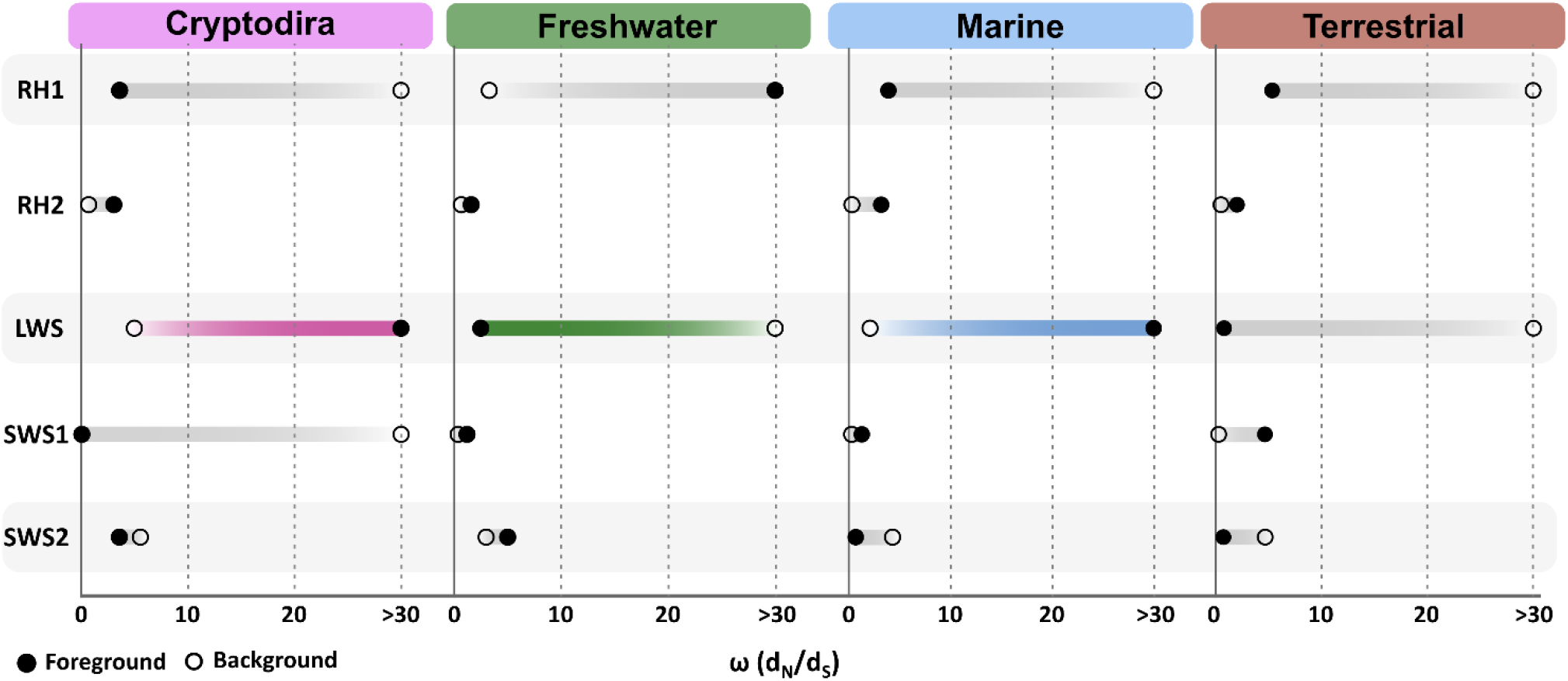
HyPhy BUSTED model results across tested turtle partitions. Estimated ω values are shown for the background (open circle) and foreground (closed circle) partitions. Partitions with evidence of positive selection are represented by coloured bars connecting the background and foreground ω values. See Supplementary File S2 for full results.

There was evidence for positive selection in a small subset of sites in marine turtle *LWS* under both topologies (ML: ω = 1186.2 [0.06%], *P* = 0.0043). To determine if the shifts in selective pressure identified by CmC are due to a relaxation of constraint or due to potential adaptive evolution, we used the RELAX model on the HYphlow platform. In the case of the marine partition, none of the visual opsins showed a significant relaxation of selection compared to non-marine turtles. In *SWS2*, the freshwater partition showed evidence for an intensification of selection whereas non-freshwater turtles showed a relaxation of selection. Interestingly we also saw evidence for an intensification of selection in Pleurodiran *SWS2* and *RH2*, but no significant relaxation of selection in Cryptodiran *SWS2* or *RH2* (Supplementary File S2).

### Variation in turtle visual opsins at known spectral tuning and kinetic rate sites

Variation was observed across turtles in the five visual opsins at gene-specific sites known to influence the spectral tuning of the other vertebrate visual pigments (Table 1; Supplemental File S3). *SWS1* had variants at the greatest number of known gene-specific sites (9), and *LWS* at the least (1). Some of the specific amino acid variants found have been previously identified in other vertebrate groups (e.g., Takahashi and Yokoyama 2005; Sugawara et al. 2009; Dungan and Chang 2016; Castiglione et al. 2017; Schott et al. 2024), but others are newly identified in turtles.

**Table 1.** Variation in turtle visual opsin sequences at known spectral tuning sites. The turtle residues we identified are listed for each spectral tuning site, with variations at that site highlighted in bold text when corresponding to that specific opsin. Site numbers are based on bovine *RH1* numbering. Spectral tuning sites following Schott et al. (2024; based primarily on Hunt et al. 2001a and Yokoyama 2008).

| Site (*RH1* numbering) | Known from | Known spectral variants | Opsin |  |  |  |  |
|---|---|---|---|---|---|---|---|
|  |  |  | *RH1* | *RH2* | *LWS* | *SWS1* | *SWS2* |
| 44 | *SWS2* | M/T | M | I | M | M/T | M |
| 46 | *SWS1/2* | F/T/L | L | F | F | **L/F/V/I** | L |
| 49 | *RH2, SWS1/2* | S/F/A/V/L | L | F | I/L/F/V/A | **F/V/L** | **V/T/A/M** |
| 52 | *RH2, SWS1/2* | L/M/T/F | F | L/I | V/I | **V/T/M/A/W** | V/M/T/L |
| 83 | *RH1/2* | D/N | N | **D/N** | D | G | N |
| 86 | *RH2, SWS1* | M/T/F/S/L/Y | M | M | E | **F/V/L** | V |
| 87 | *RH1, SWS1* | V/D/C/A | **V/I** | A | T | C | V/I |
| 90 | *RH1, SWS1* | G/S/C | G | G | A | S | G |
| 91 | *SWS1/2* | V/I/S/P | F | F | S | **V/I** | **T/S/I** |
| 93 | *SWS1/2* | T/P/L/I | T | V | I | **M/T/P** | T |
| 94 | *RH1/SWS2* | A/S/C | T | T | S | V | A/G |
| 96 | *RH1* | Y/V | Y | Y/S/C | I/V/F | I/V | Y |
| 97 | *RH2, SWS1/2* | T/A/S/C | T | T | N | **A/S** | S |
| 102 | *RH1* | Y/F | Y | Y | Y | Y | Y |
| 109 | *SWS1/2* | V/A/G | G | G | L | **A/V/G** | **A/G** |
| 113 | *RH1, SWS1* | E/D | E | E | E | E | E |
| 114 | *SWS1* | A/G | G | G | G | **A/G** | G |
| 116 | *SWS1/2* | L/V/T | F | F | T | **A/L/T** | **A/T** |
| 118 | *RH1, SWS1/2* | S/T/A/G | T | T | S | S | T |
| 119 | *RH1* | L/F | L | L | V/A/T | V/A | L |
| 122 | *RH1, SWS2* | E/I/Q/M | E | Q/K | I/V | L | M |
| 124 | *RH1* | A/S/G/V | A/G | A | A/G | T | S |
| 132 | *RH1* | A/S/V | A | A | S | A | A |
| 164 | *RH1, RH2, LWS* | S/A | A | **A/S** | **S/A** | G/S | G |
| 181 | *LWS* | H/Y | E | E | H | E | E |
| 194 | *RH1* | P/R | L | L/H | G | V | T/S |
| 195 | *RH1* | N/A | K | N/S | S/N | G | N/S/G |
| 207 | *RH1, RH2/SWS2* | M/L | M | M | L | L | L |
| 208 | *RH1* | F/Y | F | F | M | F | F |
| 211 | *RH1* | H/C | H | H | C | G/C | C |
| 261 | *RH1, SWS2, LWS* | F/Y | F | F | Y | F/Y | F |
| 265 | *RH1, SWS2* | W/Y | W | W | W | W/Y | W |
| 269 | *SWS2, LWS* | A/S/T | A | A | T | A | A |
| 292 | *RH1/2, LWS, SWS2* | A/S | A/S | A | A | A | **A/S** |
| 295 | *RH1* | A/S | A | S | A/S | S | S |
| 299 | *RH1* | A/S | **A/S** | S | T | C | T |
| 300 | *RH1* | I/T/L | I | L | I | V/I | V |
| 317 | *RH1* | A/I/M/T | M | M | I/L | I | M/L |

We found variants at three previously known spectral tuning sites in turtle *RH1* (87, 292, 299; Table 1). Site 87 was conserved as V in most sampled species, but four turtles possessed I. While V87I is not one of the known spectral tuning variants (Table 1), this substitution was also identified in frogs with evidence for positive selection (Schott et al., 2024). The effect of this variation is unknown but likely to be minor due to the similarity between the amino acid properties. At site 292, we found the spectral tuning variant S292 in *Vijayachelys sylvatica*, while A292 is conserved in all other turtles. A switch of A292S has been linked to decreases in λ_max_ of up to 10 nm in *RH1* across multiple vertebrate species, mostly associated with differences in light availability across water depths (Hunt et al., 2001; Yokoyama, 2008; Sugawara et al., 2009; Ricci et al., 2022). At site 299 variation was found between A and S, which has been shown to have a small spectral tuning effect and to affect retinal release rate in mammals (S299A, 2 nm blue shift; Dungan and Chang 2016) and was also reported to cause a small spectral shift in cichlids (Ricci et al. 2022). S299 was found in two of the pleurodiran turtles, *P. castaneus* and *P. subrufa*, whereas the remaining species were conserved for A299. The presence of A299 has been linked to a blue-shift in deep-diving fish species (Ricci et al., 2022). N83 has been suggested to be associated with both a blue shift in λ_max_ and dim-light adaptation through altered kinetic rates (Hunt et al., 2001a; Sugawara et al. 2009; Luk et al., 2016; Ricci et al., 2022). All turtle species analyzed in this dataset are primarily diurnal and are conserved for N83. Other *RH1* sites known to affect kinetic rates, such as 59, 119, 122, 124, 261, 288 (Castiglione et al. 2017; Dungan and Chang 2016; Van Nynatten et al., 2021) were conserved in both groups of turtles, as were other known tuning sites (Table 1).

*RH2* varied at two known spectral tuning sites in turtles: 83 (D/N) and 164 (A/S). At site 83 one marine cryptodiran (*Dermochelys coriacea*) possessed N, while the rest of the turtles had D83. The substitution D83N has no spectral tuning effect in chameleon *RH2* in isolation, but when combined with changes at two other sites (M86T and T97A), this caused a -15 nm shift (Takenaka and Yokoyama 2007); however, turtles are conserved as M86 and T97. The effect of this site in *RH2* on kinetic rates is likely similar to that in *RH1* but has not been studied. At site 164 turtles showed variation between A and S. While variation between A and S at 164 in *RH1* and *LWS* can have larger effects on spectral tuning, the effect of S164A was found to be very minor (-1 nm) when mutated in chameleon *RH2* (Takenaka and Yokoyama 2007).

In *SWS1*, ten spectral tuning sites with substitutions were identified: 46, 49, 52, 86, 91, 93, 97, 109, 114, and 116. Studies show that the combination of residues F46, F49, T52, F86, S90, V91, T93, A97, V109, E113, A114, L116, and S118, generates an *SWS1* with λ_max_ at 358 nm in the UV (Parry et al. 2004; Corredor et al. 2022). Cryptodiran turtles (with one exception) possess the variants F86, S90, M/T93, which are expected to result in a UV-sensitive *SWS1* pigment (Fasick et al., 2002; Hauser et al 2014; Corredor et al. 2022). In contrast, the pleurodirans in our dataset show the variants V86 and P93 which are expected to result in violet sensitivity in *SWS1* (Emerling, 2016) and are conserved for S90 (Table 2). Based on comparisons with experimentally characterized *SWS1* pigments, we predict a λ_max_ exceeding 400 nm, indicative of violet sensitivity (Table 2).

**Table 2.** Critical amino acid changes that shift λ_max_ in turtle SWS1 as seen in other vertebrates. Site numbers are based on bovine *RH1* numbering. Summarized from Hauser et al (2017).

| **Site** | **86** | **90** | **93** | **λ<sub>max</sub> (nm)** | **Shift from WT (nm)** | **References** |
|---|---|---|---|---|---|---|
| **Pleurodira** | V | S | P | Predicted >400 nm | N/A | N/A |
| **Cryptodira** | F | S | M/T | 356–358 | N/A | Loew & Govordovskii, 2001;<br>Corredor et al., 2022 |
| **Aye-aye (WT)** | F | S | P | 406 | N/A | Carvalho et al., 2012 |
| **Aye-aye P93T** | F | S | T | 371 | -35 | Carvalho et al., 2012 |
| **Goldfish (WT)** | F | S | Q | 358 | N/A | Cowing et al., 2002 |
| **Goldfish F86V** | V | S | Q | 359 | 1 | Cowing et al., 2002 |
| **Guinea pig (WT)** | V | S | A | 420 |  | Parry et al., 2004 |
| **Guinea pig V86F** | F | S | A | 367 | -53 | Parry et al., 2004 |

Variation at six known *SWS2* spectral tuning sites was found in our sample of turtles: 49, 52, 91, 109, 116, and 292 (Table 1). Most turtles have V49, but four cryptodiran (Cheloniidae) species possess T and two cryptodirans (Chelydridae) possess A. A I49V substitution was shown to cause only a small λ_max_ shift, if any, in salamander and chameleon *SWS2*, but an I49A substitution was shown to cause a more significant shift in chameleons (∼12 nm) (Yokoyama and Tada, 2003). The remaining substitution we observed at site 49, I49T, has not been investigated thus far. At site 52 we found the variants M, V, T, and L, but previous work has only shown a tuning effect of V52I/G/F/A (Yokoyama & Tada, 2003), thus the presence of these variants and the potential effect on *SWS2* sensitivity at this site is not clear. At site 91 we found the variants T, S, and I. Takahashi & Ebrey (2003), demonstrated that the substitution P91S caused a ∼10nm shift in the sensitivity of *SWS2* in newts. At site 109, we noted two different mutations, A and G, both of which have been shown to cause shifts in *SWS2* in bluefin killifish via mutagenesis, however this shift was demonstrated in conjunction with other spectral tuning sites (Yokoyama, 2008). Most turtles have A109 with the presence of G at this site restricted to tortoises (except for *Gopherus*). We recovered variations in site 116, where most turtles show A and the presence of T is restricted to two species of marine turtles and tortoises. The substitution T116L has been linked to an increase in λ_max_ in several teleosts but like site 91, these changes were found in conjunction with substitutions at other sites (Yokoyama, 2008). Finally, we noted the presence of both S and A at site 292 in turtles. Most turtles possess S292, but all Pleurodirans, and some species of softshell and marine turtles possess A292. A292S was found to shift the λ_max_ of newt (*Cynops pyrrhogaster*) by -8 nm (Takahashi & Ebrey, 2003; Yokoyama and Tada. 2003).

## Discussion

In this study, we produced the first eye transcriptome from a pleurodiran turtle and conducted analyses of the molecular evolution of turtle visual opsin genes from 59 species representing all families in both suborders (Pleurodira and Cryptodira). We used this dataset to investigate the presence and functionality of the five visual opsin genes across both suborders of turtles, and in comparison to other reptiles. We tested for differences in selection based on hypotheses of phylogenetic divergence or habitat adaptation and investigated potential spectral tuning shifts between and within suborders. All five visual opsin genes were recovered, either completely or in part, for the majority of turtle species in our dataset, supporting the long-term maintenance of photopic vision despite the inferred nocturnal ancestry of turtles. This includes expression of all five genes in the eye of the pleurodiran species *Pelusios castaneus* and presence in the eight pleurodiran genomes. The only absent gene was *SWS2* from *Gopherus evgoodei*, which we tentatively attribute to genome assembly issues rather than gene loss. We found ecological differences to be the stronger driver of differences in selective pressure across the visual opsins, more so than the deep phylogenetic divergence between the two suborders. Specifically, marine lineages best explained patterns of selection in three turtle visual opsins, while terrestrial and freshwater ecologies best explained patterns in *SWS1* and *SWS2*, respectively. Finally, we provide evidence for shifts in known spectral tuning sites in Pleurodiran *SWS1* that suggest a violet-sensitive *SWS1* pigment rather than a UV-sensitive one, which we predicted as being maintained across all sampled cryptodirans. These findings are discussed below in conjunction with our current understanding of visual opsin gene diversity, expression, function, and evolution across the vertebrate tree of life.

### Achelosaurian Visual Opsins are Remarkably Conserved Compared to Other Ancient Tetrapods

Visual opsins in turtles are under strong purifying selection, which is unsurprising considering their functional importance in the visual system. However, we did note that these ω values signalled considerably high conservation in turtle visual opsins, particularly the cone opsins, when compared to other ancient tetrapod lineages (e.g., frogs, ω = 0.096–0.113 (Schott et al., 2024); mammals (broadly) ω = 0.1-0.15 (cone opsins) & 0.04 (*RH1*) (Gai et al., 2023); bats, ω = 0.079-0.083 (Gutierrez et al., 2018)) but were comparable to previously documented reptilian M0 values (e.g. ω = 0.048–0.128 (Schott et al., 2018)). We wanted to determine if this pattern was unique to turtles by comparing overall selective constraint to other reptilian lineages. Turtle visual opsins appear to be under greater selective constraint overall compared to other reptiles for *RH2* (ω_Turtle_ = 0.056 vs ω_Reptile_ = 0.064)*, LWS* (ω_Turtle_ = 0.055 vs ω_Reptile_ = 0.073), and *SWS2* (ω_Turtle_ = 0.048 vs ω_Reptile_ = 0.060), but are under relaxed constraint for *RH1* (ω_Turtle_ = 0.140 vs ω_Reptile_ = 0.103) and *SWS1* (ω_Turtle_ = 0.047 vs ω_Reptile_ = 0.038). This partly supports one of our initial hypotheses, but the relaxed constraint on *RH1* and *SWS1* is surprising. *RH1* (rod opsin) is found in the rod photoreceptors of vertebrate retinas and is involved in scotopic (dim-light) vision (Nilsson, 2021). Rhodopsin has been the primary focus of many visual gene studies because of the documented relationship between adaptations to differing habitats and ecologies, and the availability of sequences making broad-scale studies possible (Hofmann & Lamb, 2023; Cobo et al., 2025). Due to the importance of *RH1* in dim-light environments, multiple studies have shown evidence of adaptations to differences, particularly in aquatic environments including evidence for spectral tuning (e.g. Yamaguchi et al., 2020; Ricci et al., 2022; Castiglione et al., 2023; Guo et al., 2023; Van Nynatten et al., 2024; Macpherson et al., 2024,2026) and duplication (Musilova et al., 2019). However, *RH1* across vertebrates is still remarkably conserved (Hofmann & Lamb, 2023). We did not find evidence for positive selection in *RH1* across turtles, but the significant relaxation of selective pressure may suggest variation in *RH1* is tolerated to accommodate differences or perhaps a decrease in importance given the long history of diurnality in turtles.

### No evidence of a nocturnal bottleneck in turtles

The last common ancestor of vertebrates is inferred to have five visual opsins (*RH1, RH2, LWS, SWS1,* and *SWS2*), but the retention of these genes varies extensively across extant vertebrate linages (Hagen et al., 2023). Limited visual opsin loss has been documented for ray-finned fishes (Carleton et al., 2020) and frogs (Schott et al., 2024), but extant lineages of Reptilia and Mammalia are noted as having multiple losses (Borges et al., 2018; Schott et al., 2018; Gemmell et al., 2020; Guo et al., 2023). Within Lepidosauria, visual opsin losses have been noted for geckos (*RH1, SWS2*; Schott et al., 2019a) and snakes (*RH2, SWS2*; Castoe et al., 2013), and within Archelosauria losses have been noted for crocodilians (*RH2, SWS1*; Emerling, 2016). However, another major archelosaurian lineage, birds, ancestrally maintains all five visual opsins (Borges et al., 2015). The loss or maintenance of visual opsins is often linked to the ancestral photic environment, whereby losses are expected to be more frequent in ancestrally nocturnal lineages (Lamb & Hunt, 2018; Gemmell et al., 2020). This is thought to explain in part the losses found in crocodilians but also the lack of losses in birds (Emerling, 2016, Zhang et al., 2014). Turtles, on the other hand, are inferred to be ancestrally nocturnal, suggesting the potential for visual gene loss (Anderson & Wiens, 2017). Unlike other ancestrally nocturnal tetrapod lineages, previous studies have demonstrated the maintenance of all five visual opsins across cryptodiran turtles (e.g., Horch et al., 2008; Emerling, 2016; Corredor et al., 2022), but the maintenance of these visual opsins was based on limited genomic data and was not confirmed for Pleurodirans. In concordance with previous studies, we found evidence for the maintenance of all five visual opsins broadly across Cryptodira and Pleurodira, suggesting that tetrachromatic vision is maintained across turtles. Emerling (2016) noted the possibility of gene loss or inactivation of *SWS1* in *Apalone spinifera* and both *SWS1* and *SWS2* in *Pelodiscus sinensis.* We were able to recover complete coding sequences of *SWS1* in *A. spinifera,* and *SWS1* and *SWS2* in *P. sinensis,* in contrast with Emerling’s (2016) results based on earlier genome assemblies. We did not however recover all five opsins across all genomes analyzed. In particular, we were unable to recover complete or partial sequences for *LWS* and *SWS2* for all species with *SWS2* being undetectable in the *G. evgoodei* genome assembly. In vertebrates, *LWS* and *SWS2* neighbor one another and microsynteny analyses for *G. evgoodei* and the closely related *G. flavomarginatus* revealed the presence of conserved genes flanking the two opsins in *G. evgoodei.* However, we found a discontinuity in the whole-region alignment, ambiguous sequences following the partial *LWS* locus in *G. evgoodei* and differences in gene content between the opsins. This, combined with partial and incomplete visual opsins across turtles being restricted to this specific locus is more consistent with incomplete sequencing and assembly than gene inactivation and loss. Previous studies have noted the genomes of birds are GC-rich, a characteristic known to cause difficulties in assemblies (Botero-Castro et al., 2017; Bornelöv et al., 2017; Beauclair et al., 2019; Peona et al., 2021). Additionally, crocodilians are expected to maintain three to four cone types, but the recovery of cone opsins suggests otherwise (Emerling, 2016). Synteny analyses for crocodilian opsins have highlighted genome assembly issues, rather than gene loss (Emerling, 2016). Perhaps a shared characteristic between archosaurs has created difficulties in genome assemblies. Overall, our results support a hypothesis of the continued importance of photopic vision where the ancestral opsin gene repertoire was maintained despite a possible nocturnal ancestry of turtles.

The eye transcriptome of *Pelusios castaneus* provides additional evidence that cone-mediated vision is an important component of the turtle visual system and more specifically the pleurodiran visual system. Both *LWS* and *RH1* were highly abundant, with *LWS* the most abundant individual visual opsin, while the four cone opsins were collectively more abundant than *RH1*. Cone-associated phototransduction genes were also generally more abundant than their rod-associated counterparts, although expression of *GRK7* and *GRK1* was similar. This pattern broadly resembles the complex cone systems documented in cryptodiran turtles through anatomical, physiological, and microspectrophotometric studies (Baylor & Hodgkin, 1973; Kolb & Jones, 1982; Loew & Govardovskii, 2001; Mathger et al., 2007; Grötzner et al., 2020). In *Trachemys scripta*, cones comprise most retinal photoreceptors, with L cones the most numerous cone class, followed by double and M cones, whereas rods account for a smaller proportion of the photoreceptor population (Grötzner et al., 2020). The high abundance of *RH1* and several rod-associated phototransduction transcripts nevertheless indicates a substantial rod component, consistent with the presence and physiological activity of rods documented in other cryptodiran turtles (Baylor & Hodgkin, 1973; Kolb & Jones, 1982). Because opsin transcript abundance may vary independently of photoreceptor number, these expression estimates cannot be interpreted as direct measures of rod and cone proportions. Nevertheless, the concordant abundance of multiple cone opsins and cone-associated phototransduction genes is consistent with a substantial cone component, while the high abundance of *RH1* and rod-associated transcripts indicates continued investment in rod-mediated vision. Given the deep divergence and contrasting evolutionary histories of Pleurodira and Cryptodira, these similarities raise the possibility that a complex cone system is a deeply retained feature of turtle vision.

Despite the long independent evolutionary histories and contrasting ecological diversification of Pleurodira and Cryptodira, available data provide little evidence for a major reduction of the cone system in either lineage associated with the inferred nocturnal ancestry of turtles. This contrasts with several other vertebrate lineages in which ancestral shifts towards dim-light environments have been associated with losses of visual genes and substantial restructuring of photoreceptor systems, including mammals, snakes, geckos, and crocodilians (Emerling, 2017; Borges et al., 2018; Schott et al., 2018, 2019; Pinto et al., 2019; Gower et al., 2022). In snakes, for example, *RH2* and *SWS2* were lost along the snake stem, although the ancestral snake likely retained a rod-dominated duplex retina and much of the molecular machinery underlying both rod and cone phototransduction (Schott et al., 2018; Gower et al., 2022). Crocodilians provide the closest phylogenetic comparison to turtles and retain only two intact ancestral cone-opsin classes, *SWS2* and *LWS*, whereas *SWS1* was lost and *RH2* pseudogenized, alongside losses of several other light-associated genes (Emerling, 2017). Their retinas are rod-dominated and possess multiple specialisations for dim-light vision, including a tapetum lucidum, although cones remain sufficiently diverse to support two or potentially three photopic spectral channels (Sillman et al., 1991; Nagloo et al., 2016; Karl et al., 2018). In comparison, the retention and substantial expression of all four cone-opsin classes in the pleurodiran *P. castaneus*, together with the complex cone systems documented in cryptodirans, suggests that inferred ancestral nocturnality did not produce an equally persistent reduction of the turtle cone system.

The evolutionary consequences of nocturnality are not uniform across reptiles, with tuatara providing a particularly striking counterexample to the patterns of visual-system reduction observed in other lineages with dim-light ancestry. Despite extensive dim-light adaptations, tuatara retain all five ancestral vertebrate visual opsins and nearly the complete ancestral complement of visual genes (Gemmell et al., 2020). More broadly, reptilian retinas range from essentially pure-cone to rod-dominated systems, reflecting considerable evolutionary flexibility associated with ecology and activity pattern (Katti et al., 2019). Lizards can retain and express all four ancestral cone-opsin classes (Yewers et al., 2015; Schott et al., 2019), while birds likewise generally retain complex cone systems with four spectral classes of single cones in addition to double cones and rods (Seifert et al., 2020). Together, these comparisons suggest that the effects of ancestral nocturnality on vertebrate visual systems are highly lineage dependent. In turtles, the retention of a complex cone-opsin repertoire across Pleurodira and Cryptodira, combined with evidence for substantial cone-mediated vision in both lineages, is more compatible with long-term maintenance of photopic capacity than with persistent reduction following a prolonged nocturnal bottleneck, although subsequent restoration or expansion of cone-mediated vision cannot be excluded.

### Evidence for the use of A_2_-Based Chromophores in the *Pelusios castaneus* Eye

Our recovery of substantial *CYP27C1* expression in the eye of *Pelusios castaneus*, at levels comparable to several genes involved in visual function, provides molecular evidence for an active A₁-to-A₂ conversion pathway and supports the use of A₂-based visual pigments in pleurodiran turtles. The functional link between CYP27C1 and A₂ production is well established, with experimental manipulation demonstrating that CYP27C1 is sufficient for A₁-to-A₂ conversion and required for A₂ production and its associated red shift in photoreceptor sensitivity (Enright et al., 2015; Corbo, 2021). In southern leopard frogs, differential eye transcriptome expression of *CYP27C1* corresponded with red-shifted photoreceptor spectral sensitivity in tadpoles relative to juveniles, measured using microspectrophotometry (MSP; Schott et al., 2022). Most directly relevant to turtles, A₂-based visual pigments in the cryptodiran *Trachemys scripta* are supported independently by MSP and retinal *CYP27C1* expression (Liebman & Granda, 1971; Loew & Govardovskii, 2001; Corredor et al., 2022). Liebman and Granda (1971) identified the 518-nm rod pigment of freshwater *T. scripta* as A₂-based, in contrast with an A₁-based rod pigment at ∼502 nm in the marine *Chelonia mydas*. Together, these results suggest that the capacity for A₂-based vision extends across the deep split between cryptodiran and pleurodiran turtles, while variation among cryptodirans indicates that its use differs among lineages and environments.

Across vertebrates, A₂-based visual pigments are more widespread in species inhabiting freshwater than marine or terrestrial environments, which is hypothesized to reflect spectral tuning to the red-shifted light environments that often characterize freshwater habitats (Bridges, 1972; Hauser & Chang, 2017; Corbo, 2021). The extent to which chromophore use follows habitat across turtle diversity remains an open question, however. In particular, terrestrial turtles, whose chromophore use is currently unknown, will provide an informative comparison. Indirect evidence from oil-droplet transmission in the Central Asian tortoise *Testudo horsfieldii* is consistent with A₂ use in this species. The red and yellow oil-droplet-50%-transmission wavelengths of *T. horsfieldii* closely resemble those of A₂-associated freshwater *T. scripta*, rather than the shorter-wavelength cut-offs reported for A₁-associated marine *C. mydas* (Zueva, 1982; Lipetz, 1984; Loew and Govardovskii, 2001). Although oil-droplet spectra cannot establish chromophore identity, this similarity predicts that direct measurements of *T. horsfieldii* visual pigments will reveal A₂-associated, long-wavelength-shifted sensitivities. This would not be the first evidence of A₂ use in a fully terrestrial vertebrate. Provencio et al. (1992), for example, directly identified A₂-derived chromophore in the terrestrial lizards *Anolis carolinensis* and *Podarcis sicula*, with the latter using a mixture of A₁ and A₂. More broadly, A₂ also occurs in some marine vertebrates, including California surfperches, in which MSP indicated mixed A₁/A₂ retinas, with A₂-associated pigments particularly prevalent in long-wavelength-sensitive cones (Cummings & Partridge, 2001). Chromophore composition can also vary within species rather than representing a fixed habitat-associated state (Corbo, 2021). Consequently, broader sampling of freshwater, terrestrial, and marine turtles will be necessary to determine whether chromophore use consistently tracks photic habitat or instead represents a more evolutionarily and physiologically flexible component of visual-system diversity. Such variation also complicates predictions of spectral sensitivity from opsin sequence alone, because chromophore identity can substantially alter the λ_max_ produced by the same opsin.

### Insight into Mechanisms of Oil Droplet Pigmentation from the *Pelusios castaneus* Eye Transcriptome

The *P. castaneus* eye also expressed *CYP2J19*, *BDH1L*, and *TTC39B*, genes associated with ketocarotenoid metabolism in vertebrates. *CYP2J19* is strongly expressed in the retina of the western painted turtle *Chrysemys picta* and has been implicated in the production of red ketocarotenoids that pigment the oil droplets of *LWS* cones in turtles and birds (Twyman et al., 2016). In birds, *CYP2J19* and *BDH1L* together catalyse ketocarotenoid production, while *TTC39B* enhances this activity (Toomey et al., 2022). The expression of *CYP2J19* in *P. castaneus* extends evidence for ocular expression of this pathway from cryptodiran to pleurodiran turtles, while the expression of *BDH1L* and *TTC39B* provides, to our knowledge, the first evidence that these additional components of the ketocarotenoid pathway are expressed in the turtle eye, as previously demonstrated in birds. Together, these findings are consistent with conservation of the molecular machinery underlying red cone oil-droplet pigmentation across the two major lineages of extant turtles and potentially more broadly across archelosaurian reptiles.

### Molecular Evolution of Turtle Visual Opsins is Associated with Differences in Ecology rather than Phylogeny

Given patterns of visual opsin evolution across vertebrates associated with habitat differences, we initially predicted that we would see differences in selective pressure among marine, terrestrial, and freshwater turtles. For *RH1*, *RH2*, and *LWS* the best fitting model was marine vs non-marine turtles. There is ongoing debate surrounding the ecological setting for the origin of turtles, but multiple lines of evidence point towards a terrestrial origin (Joyce, 2015). Additionally, the placement of many fossil taxa previously thought to be most closely related to extant marine lineages based on morphology is contentious and may cloud our understanding of turtle evolution (Cadena & Parham, 2015). Thus, our understanding of potential evolutionary transitions between marine, terrestrial, and freshwater ecologies across turtles creates some uncertainty regarding adaptive functional changes in turtle opsins. However, both morphology and other fossil evidence point to early turtle freshwater habitats being swampy (Joyce, 2015), likely with high presence of organic materials causing the scattering of photons similar to many modern freshwater ecosystems. This would suggest that variation in *RH1* is tolerated in marine lineages that are encountering very dim-light environments (in particular deep-diving species such as *Dermochelys coriacea* and *C. caretta*) that are also blue-shifted compared to murky freshwater environments. This partly supports our initial hypothesis for differences in selective pressure in *RH1,* where there is evidence for a release of selective constraint in turtle *RH1*. The potential for tolerating this variation can also be supported by specific reproductive and survival strategies employed by marine turtles. Both female nesting and hatchling emergence occur at night (Salmon et al., 1995; Price et al., 2018; Vandersteen et al., 2020; Simantiris et al., 2025). The importance of dim-light conditions for both nesting and hatchlings has been explored in previous studies addressing the negative impacts of artificial light on the success of both nesting and hatchling survival (Dimitriadis et al., 2018; Price et al., 2018; Chih-Hao Yen et al., 2023). Results in turtle *RH2* demonstrate overall purifying selection across the gene, but clade models show a significant difference in selective pressure between marine and non-marine turtles. Specifically, clade model results demonstrate reduced selective constraint on marine turtles relative to non-marine turtles (ω_marine_ = 0.569 *vs* ω_non-marine_ = 0.20). However, since ω remained well below one and neither BUSTED nor RELAX models detected positive selection or significant shifts in selection intensity, these results also suggest differential levels of purifying selection acting on marine vs non-marine lineages. *RH2* encodes the opsin associated with peak sensitivity in the blue-green region of the visible light spectrum, which corresponds to the dominant light wavelengths available in aquatic environments (Bowmaker, 2008; Yokoyama, 2008; Musilova & Cortesi, 2023). Within vertebrates, particularly teleost fish, previous studies have demonstrated the importance of ecological variation of aquatic light environments, including characteristics such as depth, presence of organic materials, and spectral composition (Cortesi et al., 2020; Carleton et al., 2020; Musilova & Cortesi, 2023). As mentioned previously, the environments inhabited by marine turtles vary across life history, including pelagic, coastal, reef, and in diving behaviours, each with unique spectral compositions (Bonin et al., 2006; Jaffe et al., 2011; Hochscheid, 2014; Rhodin et al., 2026). While the heterogeneous nature of these environments may weaken constraints on *RH2*, the functional importance of *RH2* in the turtle visual system may explain the reduced selective constraint on marine turtles specifically.

In *LWS* we found significant evidence that the elevated ω was due to positive selection in marine turtles, specifically. This may indicate adaptive evolution to the marine environment at a small number of sites despite overall high conservation of the gene. *LWS* is found in the cone photoreceptors of vertebrate retinas and is primarily responsible for absorbing longer “red” light wavelengths around 500–620 nm in bright-light conditions (Bowmaker, 2008). Marine environments tend to be blue shifted (greater presence of blue light wavelengths relative to others) compared to freshwater or terrestrial light environments and the impacts of this on the vertebrate visual system is well documented (Hauser & Chang, 2017; Baden, 2024). For example, previous studies tracking marine to freshwater transitions in New World Anchovies and croakers have demonstrated changes in the spectral sensitivity of *RH1* associated with the differences in light environment (Van Nynatten et al., 2015, 2021). The maximal wavelength sensitivity in *C. mydas LWS* was reported to be ∼562 nm (Liebman & Granda, 1971), which is blue shifted compared to the freshwater cryptodiran *T. scripta* (617 nm; Loew & Govordovskii, 2001). The difference in maximal sensitivity in marine turtle *LWS* is thought to be primarily driven by the use of A_1_ chromophore instead of the A_2_ chromophore used by freshwater turtles (Liebman & Granda, 1971; Horch et al., 2008; Corredor et al., 2022). Previous studies have demonstrated that the vast majority of freshwater fishes use visual pigments containing the A_2_ chromophore likely due to the increased abundance of red-light in freshwater environments (Toyama et al., 2008). Given the functional role of *LWS* in absorbing red light, the known differences in chromophore use between freshwater and marine species, and the differences in available longer light wavelengths between marine and freshwater environments this could explain the differences in maximal wavelength sensitivity between marine and freshwater turtles.

Within marine environments there is also variation in both overall light and wavelength availability (Van Nynatten et al., 2024). In clear, open-water, such as the open ocean, as depth increases the available light narrows towards blue-green (470–480 nm) (Jerlov, 1976), whereas coastal waters can become more red-shifted as blue light is absorbed or scattered by organic material (Lythgoe, 1979). Previous studies have noted shifts in visual opsin sensitivity in deep-diving/deep-dwelling species such as sharks (Hart et al., 2020; Yamaguchi et al., 2021), coelacanths (Yokoyama, 2000), whales (Dungan & Chang, 2017; Castiglione et al., 2023), and various species of both marine and freshwater ray-finned fishes (Hunt et al., 2001a; Luk et al., 2016; Musilova et al., 2019; Macpherson et al., 2024; Van Nynatten et al., 2024). Marine turtles are known to exhibit morphological adaptations and behaviours consistent with diving, however not all marine turtles dive to the same depths (Fossette et al., 2010). The leatherback turtle (*D. coriacea*) is the deepest known diver (>1000 m) moving from the epipelagic (∼0–200 m) to bathypelagic zone (∼1000–4000 m) (Fossette et al., 2010; Hochscheid, 2014). Other species such as the Loggerhead sea turtle (*C. caretta*; >600m), the green sea turtle (*Chelonia mydas*; >300 m), and the Kemp’s Ridley sea turtle (*Lepidochelys kempii*; >200 m) are known to dive from the epipelagic zone (∼0–200m) to the mesopelagic zone (∼200–1000 m) (Hochscheid, 2014). Additionally, evolutionary transitions from land to ocean are known to facilitate a series of adaptive changes (Yang et al., 2025). In marine turtles for example, these can include morphological changes (e.g., streamlined carapace, paddle shaped limbs) and adaptive changes in sensory systems including the visual system (Thewissen et al., 2007; Yang et al., 2025). Sea turtles maintain a more spherical and rigid lens compared to terrestrial or freshwater turtles, lending to a high refractive index, thought to compensate for a lack of corneal refraction (Northmore & Granda, 1991; Bartol et al., 2002). Yang et al (2025) found evidence for positive selection in a number of visual genes including those associated with eye development and evidence for specific site mutations predicted to influence gene function. These genetic changes may contribute to the distinctive visual adaptations of sea turtles relative to terrestrial and freshwater turtles (Yang et al., 2025). We found evidence for site-specific mutations at both sites known to influence spectral sensitivity and new sites in the opsin genes of marine turtles (see below). Specifically, we identified variation in *LWS* at only one known spectral tuning site in turtles. We identified both A and S residues at site 164 in turtle *LWS*, but most turtles were conserved for S164 including all marine turtles in our dataset. Several previous studies have noted that a conversion of A164S caused between a +2 to +6 nm shift via *in vitro* mutagenesis in human *LWS* (Neitz et al.,1991; Merbs & Nathans., 1992; Asenjo et al., 1994). However, as mentioned earlier, the difference in spectral sensitivities between marine and freshwater turtle species in *LWS* is expected to be approximately 55 nm which cannot be explained by the presence of S164 in marine turtles on its own. The majority of this difference in spectral sensitivity between the *LWS* based visual pigments could be attributed to the use of different chromophores. Using models based on Govardovskii et al (2000), Corbo (2021) demonstrated that the difference between an *LWS* + A_1_ visual pigment and an *LWS* + A_2_ visual pigment is approximately 41 nm. The site-specific mutation identified in marine turtle *LWS* in combination with the known differences in spectral sensitivity resulting from differences in chromophore usage could explain most of the previously documented differences in spectral sensitivity between marine and freshwater turtles.

Another outcome of marine-specific morphological adaptation is an increased locomotion speed in marine species (Marvin & Lutterschmidt, 1997; Eckert, 2002). We initially hypothesized that because marine turtles were moving through a blue-shifted photic environment quickly, that changes in visual opsins would be apparent in *RH1*. We also predicted that in addition to site-specific changes that alter spectral sensitivity, we would see alterations that affect kinetic rates. While we did not find evidence of positive selection in *RH1* in marine species or turtles more broadly, there were marine-specific patterns (discussed above). Perhaps the shift between red-light dominated photic environments in freshwater and blue-light dominated marine photic environments, which are also depth dependent, may explain the patterns seen in marine turtle *RH1* and *LWS*, along with the differences in locomotion. While previous studies have explored site-specific changes that affect kinetics in *RH1* (e.g., retinal release rate (Morrow and Chang, 2015; Van Nynatten et al., 2021) these are less well understood in cone opsins (e.g., Hart et al., 2019). We noted two specific amino acid changes in marine turtle *LWS* (G155A and Y37F) that have not been explored previously. While much of the spectral tuning of marine turtle opsins is thought to be related to differences in chromophore use, signals of positive selection could reflect other functional adaptations such as changes to kinetic rates, the efficiency of activation/deactivation of the photoreceptor, or the fine tuning of functions directly associated with the chromophore.

For *SWS1*, the best-fitting model was terrestrial vs aquatic turtles. *SWS1* is also found in the cone photoreceptors of vertebrate retinas and is maximally sensitive to UV/violet light wavelengths (“ultraviolet”, λ_max_ ∼355–375 nm, or “violet”, ∼385–455 nm) (Bowmaker, 2008; Hauser et al., 2014). As mentioned previously there are significant differences in light wavelength composition and availability between environments, particularly between aquatic and terrestrial environments (Baden, 2024). Light is both scattered and absorbed strongly in water relative to air, and as a result more photons of light are available and viewing distance is increased in terrestrial environments (Baden, 2024). Transitions between terrestrial and aquatic environments have affected the evolution of various visual opsins including the short wavelength sensitive opsins in multiple vertebrate lineages (e.g. frogs (Schott et al., 2024), sea snakes (Seiko et al., 2020)), and marine mammals (Fasick et al., 1998, Fasick & Robinson, 2000; Southall et al., 2002; Levenson et al., 2006; Castiglione et al., 2023), and include examples of both gene loss and spectral tuning. Our clade model results provide evidence for a difference in selective pressure between terrestrial and aquatic turtles in *SWS1*, specifically, clade model results show terrestrial turtles under greater selective constraint compared to aquatic turtles (ω_terrestrial_ <ω_aquatic_). Additionally, there was no evidence of positive selection in either PAML or BUSTED analyses and RELAX analyses detected no significant changes in selection intensity among these groups. This pattern suggests that *SWS1* function remains highly conserved in turtles more broadly but also where terrestrial turtles demonstrate particularly strong evolutionary constraint. As mentioned *SWS1* confers UV/violet light sensitivity in vertebrates. Terrestrial turtles are predominately diurnal and are most often found in open habitats that are also associated with high irradiance of UV light (Lagarde et al., 2002; Bonin et al., 2006; Baden, 2024). In these environments visual cues are important for foraging, navigation, and social interactions, all of which may have specifics in the UV spectrum (McRae et al., 1981; Honkavaara et al., 2002; Cronin & Bok, 2016). Perhaps the increase of selective constraint in terrestrial turtles represents a stronger evolutionary conservation of the UV sensitive function of *SWS1* in this group relative to aquatic species.

For *SWS2* the best-fitting model was freshwater vs non freshwater turtles. *SWS2* is also found in the cone photoreceptors of vertebrate retinas and is maximally sensitive to blue light wavelengths (λ_max_ 400–470 nm) (Bowmaker, 2008). As mentioned previously there are significant differences in light wavelength composition and availability between environments. Much like marine environments, this wavelength availability and composition also vary across freshwater environments (Hauser et al., 2021; Costa et al., 2013). Freshwater habitats can be classified into multiple types: clearwater, with its high transparency, allows wavelengths across the visible spectrum to penetrate (including UV wavelengths); blackwater, which is high in dissolved tannins creating a domination of orange/red wavelengths; and white water, which has low transparency due to higher turbidity and increased sediment presence (Albert & Reis, 2011, Costa et al., 2013). As a result of these characteristics, both white water and black water habitats show a disproportionate attenuation and scattering of short light wavelengths (Costa et al., 2013). Previous studies in fishes have demonstrated that loss and retention (including expression) of both *SWS1* and *SWS2* are highly dependent on habitat (Allison et al., 2003; Hofmann et al., 2010; Cortesi et al., 2020; Hauser et al., 2021). In neotropical freshwater cichlids, there is evidence of spectral tuning in *SWS2a* (fishes possess a duplication of the *SWS2* genes), thought to be the primary method of tuning to short light wavelengths in lineages that have lost *SWS1* across these variable freshwater environments (Hauser et al., 2021). In Beloniformes, both short wavelength sensitive opsins (*SWS1* and *SWS2*) were found to be under strong purifying selection thought to be related the functional importance of these genes (Chau et al., 2024). Significant shifts in selection in *SWS2* were also noted between freshwater and marine Beloniformes lineages, which was thought to be related to differences in available light between these two environments (Chau et al., 2024). Most extant turtle species inhabit freshwater environments. However, we are limited by the availability of sample data and natural history information that would be needed to further investigate the effect of different freshwater habitat types more closely. Interestingly, our clade model results show freshwater turtle *SWS2* to be under greater selective constraint (ω=0.053) relative to non-freshwater turtles (ω=0.416). The significant differences in selective pressure we noted between freshwater and non-freshwater turtles were also accompanied by evidence for an intensification of selection in freshwater turtles (*k* = 1.27, *p* = 0.0081), and a relaxation of selection in non-freshwater turtles (*k* = 0.82, *p* = 0.0215). In both cases however, we found no evidence for positive selection in *SWS2* in our BUSTED analyses in our turtle only dataset, but there was evidence for positive selection in turtles compared to other reptiles (ω_turtle_ = 9.92, p = 0.03887). These results suggest that long-term shifts in selection across freshwater lineages are driven primarily by sustained changes in selection intensity, in this case an intensification of negative selection, that may be related to differences in water types. However, signals of positive selection in turtle *SWS2* relative to other reptiles may suggest previous adaptive events deep in turtle evolutionary history. Given the known variation in light wavelength availability across freshwater environments, perhaps this suggests that in freshwater environments the maintenance of a diverse opsin repertoire is conserved over evolutionary time.

### Spectral Tuning of Turtle Visual Pigments

We identified variation at spectral tuning sites in both turtle groups for each of the five visual opsins including numerous new variants that are likely to have shifted the spectral sensitivities of turtle visual pigments. Most notably, we uncovered a mutation shared by the pleurodiran *SWS1* sequences, M93P, that in combination with F86V, we inferred shifts sensitivity from UV to violet. All cryptodiran *SWS1* are measured, or inferred, to be UV sensitive (Loew & Govordovskii, 2001; Corredor et al., 2022), indicating a potentially underappreciated difference in the visual ecologies of the two suborders.

The *SWS1* pigments in both *Chrysemys picta* and *Chelonia mydas* were predicted to be ultraviolet-sensitive due to the presence of F86, S90, and M93 or T93 (Emerling, 2016), and electroretinography (ERG) measurements suggested an increase in sensitivity towards the UV and supported the presence of a well-developed photopic visual system in these two species of turtles (Levenson et al. 2004). These results were also supported by behavioral training techniques and *in vitro* expression of reconstituted opsins that confirmed the presence of a UV-sensitive photoreceptor (S-cone) in the retina of turtles (Arnold and Neumeyer, 1987; Ventura et al. 2001). Ultraviolet vision in marine turtles is also supported by both the presence of UV fluorescing oil droplets and measurements of the spectral transmission patterns of ocular media (Mäthger et al., 2007). Studies on the role of UV sensitivity in cryptodiran behaviour have focused mainly on marine turtles. Numerous investigations have affirmed that hatchling sea turtles, particularly loggerhead turtles (*Caretta caretta*), predominantly emerge during the night, irrespective of the moon’s phase or position (Kawamura et al. 2009). Notably, even when situated 50 meters away from the sea, they move seaward, although the sea is not visually discernible to them at that distance. Their primary mechanism for locating the sea is oriented toward light, as evidenced by the fact that blindfolded hatchlings struggle to find the sea (Kawamura et al. 2009). Loggerhead turtle hatchlings exhibit a deliberate avoidance of yellow and orange light, showing a strong attraction to light within the near ultraviolet to the green portion of the spectrum. This light preference is not exclusive to loggerhead turtles; other species such as green turtles (*Chelonia mydas*), olive ridley turtles (*Lepidochelys olivacea*), and hawksbill turtles (*Eretmochelys imbricata*) also demonstrate effective orientation responses to near-UV light (Witherington & Bjorndal, 1991). Given that near-UV light is more dominant over the sea compared to land, irrespective of factors such as latitude, moon phase, and sky condition, it serves as a critical visual cue for hatchlings. In contrast, light is absorbed considerably by land and vegetation, and clouds, while dimming and diffusing light, do not significantly alter its spectral composition within the 350–800 nm range (Kondratyev et al. 1969). This could explain the importance in maintaining UV sensitivity in at least marine turtles.

Our understanding of the importance of UV sensitivity in other freshwater and terrestrial cryptodirans is still not well understood. Birds like turtles maintain tetrachromatic vision and previous studies have demonstrated evidence for both UV and violet sensitivity in *SWS1* (Borges et al., 2015). The ecological role of *SWS1* in birds is thought to be broad encompassing roles associated with colour pattern recognition (Ödeen & Hǻstad, 2010), social signalling, hunting, nectar localization, and mate-choice (Burkhardt, 1989; Viitala et al., 1995; Bennett et al., 1996). Violet sensitivity is thought to be the ancestral condition of bird *SWS1*, however this is still contentious (Ödeen & Hǻstad, 2013; Borges et al., 2015). Ödeen & Hǻstad (2013) showed that across the 21 avian orders studied, shifts in *SWS1* between UV and violet sensitivity occurred at least 14 times and were attributed to specific site substitutions at sites 86, 90, and 93. The ability to discriminate between UV and visible light is thought to lead to better colour discrimination (Ödeen & Hǻstad, 2013). One explanation for a shift in absorption in birds is related to the cost of increased UV transmission as shortwave radiation which can cause photooxidation in biological tissue (Ödeen & Hǻstad, 2013). This may result in selective pressures related to the protection of the retina from the damaging effects of UV light and potentially an improvement in image quality on the retina when UV sensitivity or better colour discrimination is not more beneficial (Hunt et al., 2009). The switch to violet sensitivity in pleurodiran turtles could then be related to retinal protection however it is not clear why this would be the case in pleurodirans but not freshwater cryptodirans.

The *RH1* site (N83) is believed to be associated with adaptation to dim-light conditions (Sugawara et al., 2009). Interestingly, all turtles in our dataset exhibited the N83 variant. The ecological origins of turtles remain a contentious topic in reptile evolution. Some studies suggest a relationship with marine sauropterygians (Li et al., 2008), while others propose a terrestrial or fossorial origin (Joyce et al. 2004; Emerling, 2016; Lyson et al., 2016). The presence of N83 could be consistent with a nocturnal origin for turtles, but the maintenance of all ancestral visual opsins argues strongly against a fossorial one. The observed consistency in the N83 site across all turtles in our dataset prompts further exploration into the specificities of their visual adaptations and evolutionary history in relation to their ecological niches. Several studies have indicated the significance of D83N and A292S in causing spectral blue shifts in rhodopsin pigments found in deep-sea creatures and marine animals (Hunt et al. 2001a; Dungan and Chang 2016). This adaptation facilitates optimal photon capture in the more blue-shifted light conditions prevalent underwater, but all turtles except one species retained A292, while *Vijayachelys silvatica* possessed S292.

## Conclusions

The existing body of knowledge pertaining to the visual opsins and their spectral sensitivity in turtles has primarily focused on species within the Cryptodira suborder. The present study represents the first investigation into the photoreceptor cells and spectral sensitivities of pleurodiran turtles. We find evidence for the maintenance and expression of all five visual opsins across both suborders and that the limited evolutionary change in these genes is driven primarily by ecological differences. We also found that despite high levels of conservation in turtle visual opsins, this pattern is only unique to turtles in three of the visual opsin genes, whereas archelosaurs, together, have more conserved visual opsins than many other vertebrate lineages. Our results highlight the potential functional adaptation of pleurodiran *SWS1* from UV to violet sensitivity through evidence for substitutions at known spectral tuning sites. Future studies should measure the spectral sensitivity of the visual opsins in pleurodiran species using MSP or *in vitro* expression to determine the extent of UV or violet sensitivity of the *SWS1* opsin in this suborder of turtles and compare it to cryptodiran turtles. A review of the visual adaptation differences between the two suborders can aid in the investigation of potential reasons behind the observed differences in selection of spectral sensitivity in visual genes of the two groups. While the present study validated the expression of *CYP27C1* in pleurodirans, future studies should test for the expression of this gene in cryptodiran turtles as well. The outcomes of our research contribute significantly to enhancing our comprehension of the functional aspects of visual opsins and the adaptive mechanisms inherent in this sensory modality within pleurodiran turtles.

## Acknowledgments

This study was support by NSERC Discovery Grants to RKS and BSWC and by the National Science Foundation under the Summer Systematics Institute REU site grant number 1852276.

## Data Availability Statement

Upon publication data supporting the study will be made available including sequencing reads on NCBI SRA and the transcriptome assembly, gene alignments, phylogenies and other results files on Zenodo.

